# From gene to structure: Unraveling genomic dark matter in *Ca*. Accumulibacter

**DOI:** 10.1101/2024.05.14.594254

**Authors:** Xiaojing Xie, Xuhan Deng, Liping Chen, Jing Yuan, Hang Chen, Chaohai Wei, Chunhua Feng, Xianghui Liu, Guanglei Qiu

**Affiliations:** School of Environment and Energy, South China University of Technology, Guangzhou 510006, China; Singapore Centre for Environmental Life Sciences Engineering, Nanyang Technological University, Singapore 637551, Singapore; Guangdong Provincial Key Laboratory of Solid Wastes Pollution Control and Recycling, Guangzhou 510006, China; The Key Lab of Pollution Control and Ecosystem Restoration in Industry Clusters, Ministry of Education, Guangzhou 510006, China

**Keywords:** *Candidatus* Accumulibacter, Alphafold2, Protein structure, function annotation

## Abstract

*Candidatus* Accumulibacter is a unique and pivotal genus of polyphosphate-accumulating organisms (PAOs) prevalent in wastewater treatment plants, and plays mainstay roles in the global phosphorus cycle. Whereas, the efforts toward a complete understanding of their genetic and metabolic characteristics are largely hindered by major limitations in existing sequence-based annotation methods, leaving more than half of their protein-encoding genes unannotated. To address the challenge, we developed a comprehensive approach integrating pangenome analysis, gene-based protein structure and function prediction, and metatranscriptomic analysis, extending beyond the constraints of sequence-centric methodologies. The application to *Ca*. Accumulibacter allowed the establishment of the pan-*Ca*. Accumulibacter proteome structure database, providing references for >200,000 proteins.

Benchmarking on 28 *Ca*. Accumulibacter genomes showed major increases in the average annotation coverage from 51% to 83%. Genetic and metabolic characteristics that had eluded exploration via conventional methods were unraveled. For instance, the identification of a previously unknown phosphofructokinase gene suggests that all *Ca.* Accumulibacter encoded a complete Embden-Meyerhof-Parnas pathway. A previously defined homolog of phosphate-specific transport system accessory protein (PhoU) was actually an inorganic phosphate transport (Pit) accessory protein, regulating Pit instead of the high-affinity phosphate transport (Pst), a key to the emergence of the polyphosphate-accumulating trait of *Ca.* Accumulibacter. Additional lineage members were found encoding complete denitrification pathways. This study offers a readily usable and transferable tool for the establishment of high-coverage annotation reference databases for diverse cultured and uncultured bacteria, facilitating the exploration and understanding of genomic dark matter in the bacterial domain.

**Synopsis:** A integrated and advanced approach unraveling key genomic dark matter in *Ca*. Accumulibacter and readily applicable to diverse bacteria for customerized high-coverage annotation reference database establishment

## Introduction

The phosphors (P) cycle, shaped by human interventions to overcome P limitation, has significantly accelerated P inputs into the biosphere [1–3]. Human activities produce P-containing wastewater, which is a main driver of large-scale anthropogenic P release into aquatic ecosystems and causes disastrous consequences [4, 5]. Current mitigation strategies focus on resolving P flows from soil and crops and improving the removal and recovery of P in wastewater to sustainably manage the intricate challenges of the altered P cycle [6, 7]. Enhanced biological phosphorus removal (EBPR) is widely applied in full-scale wastewater treatment plants (WWTPs) for P removal and recovery owing to its economical and sustainable features [7, 8]. This process is mediated by a group of microorganisms, namely polyphosphate-accumulating organisms (PAOs), having an extraordinary ability to release and excessively take up phosphate under anaerobic and aerobic conditions, respectively [9, 10]. *Candidatus* Accumulibacter is a model PAOs [6, 11, 12]. To confer stable EBPR with improved P removal efficiencies, the metabolic characteristics of this group of bacteria were extensively studied [13–17]. Culture-independent high-throughput sequencing, and sequence-based comparative and systematic genomic analyses offered key approaches for their metabolic potential studies, spilling beans on their genetic secrets behind the unique traits [18–20]. Whereas, the availability, identification and functional annotation of genes in an organism is a key to genomic analyses. Despite the intensive studies, 40%-60% protein-coding genes in *Ca*. Accumulibacter remain uncharacterized with nameless functions, greatly hindering a complete elaboration of their metabolic characteristics [21, 22].

Current genome-scale functional annotation methods share the same major limitation of relying on function prediction by homology searching against public protein reference databases, such as COG, and/or KEGG orthology [21, 23]. Annotation methods based on homology are fast; however, suffer from low coverage. Any gene sequence differs from reference protein-encoding gene families is inevitably ignored and excluded in subsequent analyses. Additionally, conventional homologous annotation methods typically assume that similar sequences correspond to similar functions. However, in a biological context, similar sequences may encode proteins with distinct structures and functions, challenging the accuracy and rationality of functional annotations [24–26]. Consequently, incomplete and even prejudiced information may be obtained, imposing significant obstructions on the understanding and exploration of the metabolic characteristics of bacteria. To reduce the incompleteness of information and assess the extent of unexplored functional diversity, a comprehensive approach like pangenomics, involving all vs all genomic comparison, becomes necessary [11, 20, 27]. However, such analyses demands substantial computational resources, entails intricate subsequent analysis, nevertheless, remains powerless in handling the huge amounts of hypothetical proteins. Although efforts to dig deeper into genomic dark matter have been recently reported, there is still a lack of universally investigative tools for the comprehensive study of cultured and uncultured bacteria [28–30].

The intricate relationship between genome sequences and biological functions may be illuminated by the three-dimensional (3D) structures of proteins [31, 32]. Despite the commonly occurring gene sequences with low identities, the discovery that they can encode similar protein structures implies potential functional congruence [33, 34]. In the past decades, traditional methods for protein structure characterization and determination in newly sequenced organisms have been extremely difficult and greatly intractable [35, 36]. Recent breakthroughs, exemplified by powerful deep learning tools such as AlphaFold2, have revolutionized our ability to derive proteome-wide structures from genome sequences with high-reliability [37, 38]. In addition, deep learning methods for function prediction based on protein structure have been shown to generate highly accurate functional annotations, even for novel sequences without homologs in public databases [39–41]. These methods can predict protein function by leveraging sequence features extracted from protein language models and structures, enabling unprecedented resolutions and insights into protein functions. These advances have a transformative impact and the potential to address a critical challenge in genomics—bridging the gaps between sequence information and biological function [42–44], enabling a more intuitive and in-depth understanding of functional characteristics, surpassing the limitations of genomics alone.

In this study, combined with protein structure prediction and analysis, and comparative genomics, we developed a comprehensive approach with universal applicability to uncover previously unknown metabolic characteristics in *Ca.* Accumulibacter. Via protein structure clustering and function prediction, our result demonstrated significantly enhanced annotation coverage compared to the widely used COG or KEGG database. Alphafold2 was further utilized to predict the structures of 60 polyphosphate kinase 1 (PPK1) and 52 polyphosphate kinase 1 (PPK2) for sequence- and structure-based phylogenetic analyses, revealing the complex evolutionary process of *Ca*. Accumulibacter. Comparative analyses among *Ca*. Accumulibacter species and with *Ca*. Propionivibrio unveiled unexplorable genomic features via conventional sequence-based annotation. For instance, the successful identification of a previously unknown phosphofructokinase gene (*pfk*) suggests that all *Ca.* Accumulibacter encoded a complete Embden-Meyerhof-Parnas (EMP) pathway for glycolysis. A protein previously thought to be a homolog of phosphate-specific transport system accessory protein (PhoU) was redefined as putative inorganic phosphate transporter (Pit) accessory protein (Pap), and would have been involved in the regulation of Pit rather than the high-affinity phosphate transporter (Pst), a key to the development of the polyphosphate-accumulating trait of *Ca.* Accumulibacter. Structural analysis further revealed additionally *Ca*. Accumulibacter lineage members encoding complete denitrification pathways, undiscoverable via sequence-based annotation. Integrating pangenomic analysis and sequence- and structure-based annotation, an annotated reference database was built for *Ca*. Accumulibacter, achieving major improvements in extending the range of sequence-based annotations. This study illustrated a successful integration of protein structure and functional prediction, and protein-protein interactions to gain innovative metabolic insights, revealing unexplored and diverse functional spaces in *Ca*. Accumulibacter. The method developed in this work is readily applicable and transferable to other cultured and uncultured bacteria, for the exploration of genomic and functional dark matter in the bacterial domain.

## Material and Methods

### Enrichment, metagenomic and metatranscriptomic analyses of *Ca*. Accumulibacter

A sequential batch reactor (SBR) was used for the enrichment of *Ca*. Accumulibacter. The SBR has a working volume of 4.5L and was inoculated with activated sludge from a WWTP in Guangzhou, China [45, 46]. The SBR was operated with 6 h cycles, feeding on a synthetic medium (detailed in the Supplementary information) with acetate as a sole carbon source. The obtained enrichment cultures (i.e., *Ca.* Accumulibacter cognatus SCUT-2 and SCUT-3) were subjected to anaerobic-aerobic full-cycle studies and metatranscriptomic analyses to validate the metabolic potentials obtained in this study. Details on the reactor operation, anaerobic-aerobic full-cycle study, sample collection, metagenomic and metatranscriptomic analyses were documented in the Supporting Information (SI). The raw reads and draft genomes obtained have been uploaded to the NCBI GenBank database under the BioProject No. PRJNA807832, No. PRJNA994326 and No. PRJNA771771.

### Data acquisition and evaluation

A total of 89 high-quality *Ca*. Accumulibacter metagenome-assembled genomes (MAGs) were obtained including 11 recovered from our laboratory-scale enrichment reactors and 78 from the NCBI database [47]. CheckM2 was used to assess the genome integrity and degree of contaminations [48]. After quality assessment, 47 high-quality *Ca*. Accumulibacter genomes (completeness >95%, contamination <6%) and 12 moderate-quality *Ca*. Accumulibacter genomes were selected. In addition, 2 *Ca*. Dechloromonas genomes (both are PAOs) [49], and 1 *Ca*. Propionivibrio aalborgensis genome (a GAO genome) [50] were included in the analysis as outgroups and to facilitate the comparison between PAOs and GAOs. The GenBank or RefSeq assembly accession and additional details about the qualities of these genomes can be found in the SI Table S2.

### Protein structure prediction

Three-dimensional (3D) structure of 60,198 protein sequences of 17 *Ca*. Accumulibacter genomes, and 3,602 protein sequences of 1 *Ca*. Propionivibrio aalborgensis genome were obtained from the Alphafold database (SI Table S3) [51]. The confidence in the accuracy of the predicted structures was assessed through the predicted local distance difference test (pLDDT). Residues with pLDDT scores ≥70 were classified as high confidence and pLDDT scores <50 were considered as low confidence. rafm was used to calculate the pLDDt scores [52]. In addition. Alphafold2 was used to predict the 3D structure of proteins in key (carbon, phosphorus and nitrogen) metabolic pathways of *Ca*. Accumulibacter. For each prediction, five models were generated, ranked from 1 to 5 with 1 as the highest pLDDT score. The results obtained using five different models are generally consistent, the model with the highest pLDDT was used for further analysis. For metabolic insights, we focused on structures with pLDDT scores > 70, ensuring result accuracy. Pymol was used to visualize the protein structures.

### Clustering based on the sequence and structure of the protein

For orthologous gene cluster identification based on the protein sequences, all vs all BLAST of each genome was performed using Orthofinder 2.5.4 with parameters -evalue 1e-5, -seg yes, -soft_masking true, -use_sw_tback. The obtained results were filtered to a query coverage of 75% and a percent identity of 70% [20]. Orthologous gene clusters were identified using MCL version 14-137 with an inflation value of 1.1 [53]. Foldseek was used for structural clustering to find proteins with similar structures with parameters -evalue 1e-4- covered 0.9 [54].

### Structure- and sequence-based protein function prediction

To elucidate the metabolic potential of *Ca*. Accumulibacter, KEGG and DeepFRI were used for functional annotation based on protein sequence and structures, respectively [21, 39]. The quality of DeepFRI predictions were defined as high quality with scores>0.5, and standard quality with scores >0.2. To confirm the annotation viability, DeepFRI annotations with a threshold above 0.2 were considered to be significant. We further stratified annotations into three categories: (I) successful annotations with both methods; (II) DeepFRI unique (genes successfully annotated with DeepFRI only); and (III) KEGG unique (genes successfully annotated with KEGG only). STRING was used to identify interacting genes/proteins [55]. Cytoscape was used for PPI network visualization [56]. To simulate possible protein interactions, Hdock was used to calculate protein-protein docking based on their structures [57]. Additionally, the outcomes obtained from DeepFRI and KEGG were incorporated along with the sequence clustering results. These data were used to supplement the functional annotation of the pan *Ca*. Accumulibacter genome. Subsequently, A refined annotation database was established, achieving major improvements in the annotation efficiency of *Ca*. Accumulibacter.

### Phylogenetic analyses

Phylogenetic analyses of *Ca.* Accumulibacter were performed based on the pangenome, PPK1 gene sequences and structures, and PPK2 gene sequences and structures, respectively. For the phylogenetic analysis based on pangenome, an STAG algorithm was used. Rootless trees were constructed by using all genes instead of single-copy genes [58]. The STRIDE algorithm was used to infer rooted trees from the rootless trees [59]. Maximum likelihood trees of PPK1 and PPK2 genes were constructed by using iqtree2 with parameter -b 100 -T 30, using model GTR+F+I+G4 and TIM3+F+I+R3, respectively [60]. foldtree was used to construct phylogenetic trees based on the 3D structure of PPK1 and PPK2 to analyze their evolutionary relationships [43]. Landscaping of the phylogenetic trees was achieved using Chiplot [61].

## Results and Discussion

### Three-dimensional structure prediction and annotation

The 3D structure of proteins encoded by 127 genes, including 60 *ppk*1, 52 *ppk*2, and 3 denitrification genes (encoding the small and large subunits of nitric oxide reductase NorC, NorB, and nitrous oxide reductase NosZ), 3 *pho*U, 1 *pst*S, 4 *pho*U homologs, and 4 *pit*, were predicted with high confidence levels (exceeding 70) (SI Figure S2) [37]. Simultaneously, we analyzed 60,198 gene-encoding protein structures from 11 *Ca.* Accumulibacter datasets, attributing to 17 *Ca.* Accumulibacter genomes from five clades (SI Figure S1). The average number of residues possessed by those proteins was 298.4, maximized at 1280, and minimized at 18 (Figure 1C). The average pLDDT score of those *Ca.* Accumuibacter encoding protein structures (60,198) was 83.9, with 87.8% of them (52,853 of 60,198) having pLDDT scores >70, indicating that the predicted structures are highly confident (Figure 1B).

**Figure 1.**
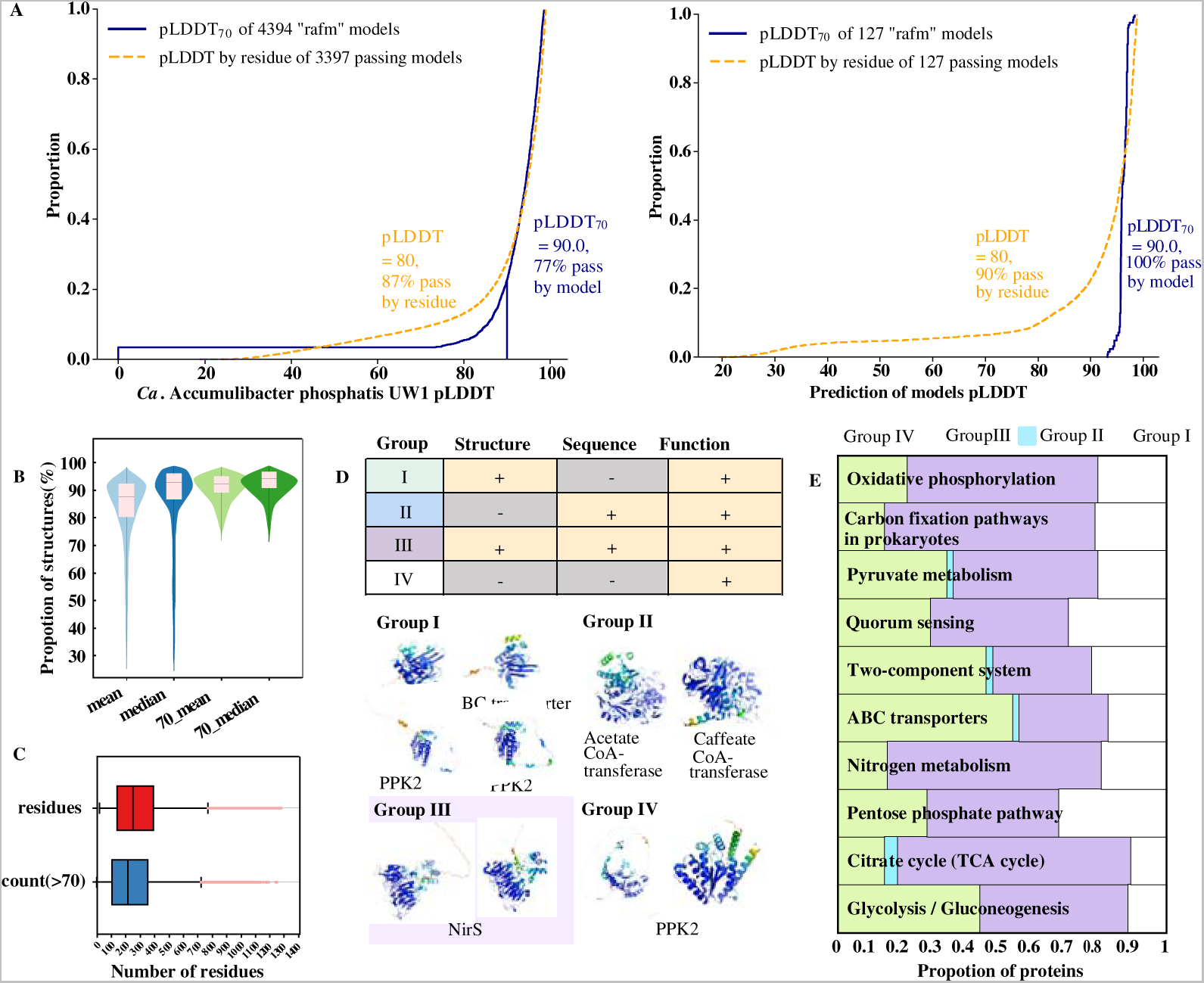
(A) The pLDDT confidence scores of proteins in *Ca*. Accumulibacter phosphatis UW1 and 127 predicted protein structures. (B) The mean and median of residue pLDDT scores of 60,189 proteins; and the mean and median of residues with pLDDT score >70. (C) A box plot showing the distribution of number of residues and those with a pLDDT score >70 in each model. (D) 3D structure prediction and annotation. Classifications of proteins are based on the similarities of their structures, sequences and functions between protein pairs of *Ca*. Accumulibacter similis SCELSE-1 and *Ca*. Propionivibrio aalborgensis S1. Proteins with the same or similar functions were classified into four groups. Representative cases of protein comparison between *Ca*. Accumulibacter (left) and *Ca*. Propionivibrio (right) with the same/similar functions in three groups. (E) The proportion of protein classification in each metabolic pathway of SCELSE-1.

The molecular function (MF) and enzyme commission (EC) descriptions of all proteins (60,198) were predicted using DeepFRI, generating probability scores within the range of 0 to 1 (SI Spreadsheet 2). *Ca.* Accumulibacter similis SCELSE-1 recovered from our lab-scale reactor [45], was selected as a representative genome for annotation comparison. SCELSE-1 comprises 3,559 genes with 1,992 of them (56.0%) were successfully annotated based on KEGG. The functions of 3,299 gene-encoded proteins were predicted using DeepFRI. After excluding predictions with confidence scores below 0.2/0.5, 3149/2065 proteins were successfully annotated. Notably, 1214/646 of them were unannotated with the KEGG database, suggesting major improvement in the annotation coverage. 900 protein-protein interaction (PPI) networks were observed for SCELSE-1, of which, 665 involved hypothetical proteins. The validity of the predicted PPI was demonstrated by known networks, such as phosphate metabolism networks (Figure 2C). The integration of PPI predictions with DeepFRI unearthed novel features within the network involving hypothetical proteins. For instance, for the PPI network involving FAZ92_02561, FAZ92_00531, FAZ92_00791, FAZ92_01419, and FAZ92_01104, the proteins FAZ92_02561 and FAZ92_00531, identified as PhaC based on KEGG, were pivotal enzymes in polyhydroxyalkanoate (PHA) biosynthesis. DeepFri annotation suggested that FAZ92_00791 played a regulatory role towards those PhaC proteins. By examining the metatranscriptome of *Ca.* Accumulibacter cognatus SCUT-3 (SI Spreadsheet 3), genes encoding FAZ92_00791 and FAZ92_00531 (PhaC) showed the same transcriptional trend, both of which were highly transcribed throughout the anaerobic-aerobic EBPR cycle, suggesting a positive regulatory role of FAZ92_00791 towards PhaC. The PHA granules consist of a polyester core, surrounded by a boundary layer with embedded or attached proteins including the PHA synthase and phasins, depolymerizing enzymes, and regulatory proteins [62]. FAZ92_01419, with structural molecule activity and lipid-binding ability based on its structure (Figure 2D), would have contributed to the structural integrity of PHA granules, which was also highly transcribed throughout the anaerobic-aerobic EBPR cycle for *Ca.* Accumulibacter cognatus SCUT-3 (SI Spreadsheet 3). Collaboratively, FAZ92_01419 and FAZ92_00791 may have regulated the formation and maintained structural stability of PHA granules in *Ca.* Accumulibacter. Meanwhile, FAZ92_01104, possessing transport activity based on DeepFRI, exhibited low transcription in *Ca.* Accumulibacter cognatus SCUT-3 (a maximum RPKM of 132, SI Spreadsheet 3), suggesting limited involvement in actual functions.

**Figure 2.**
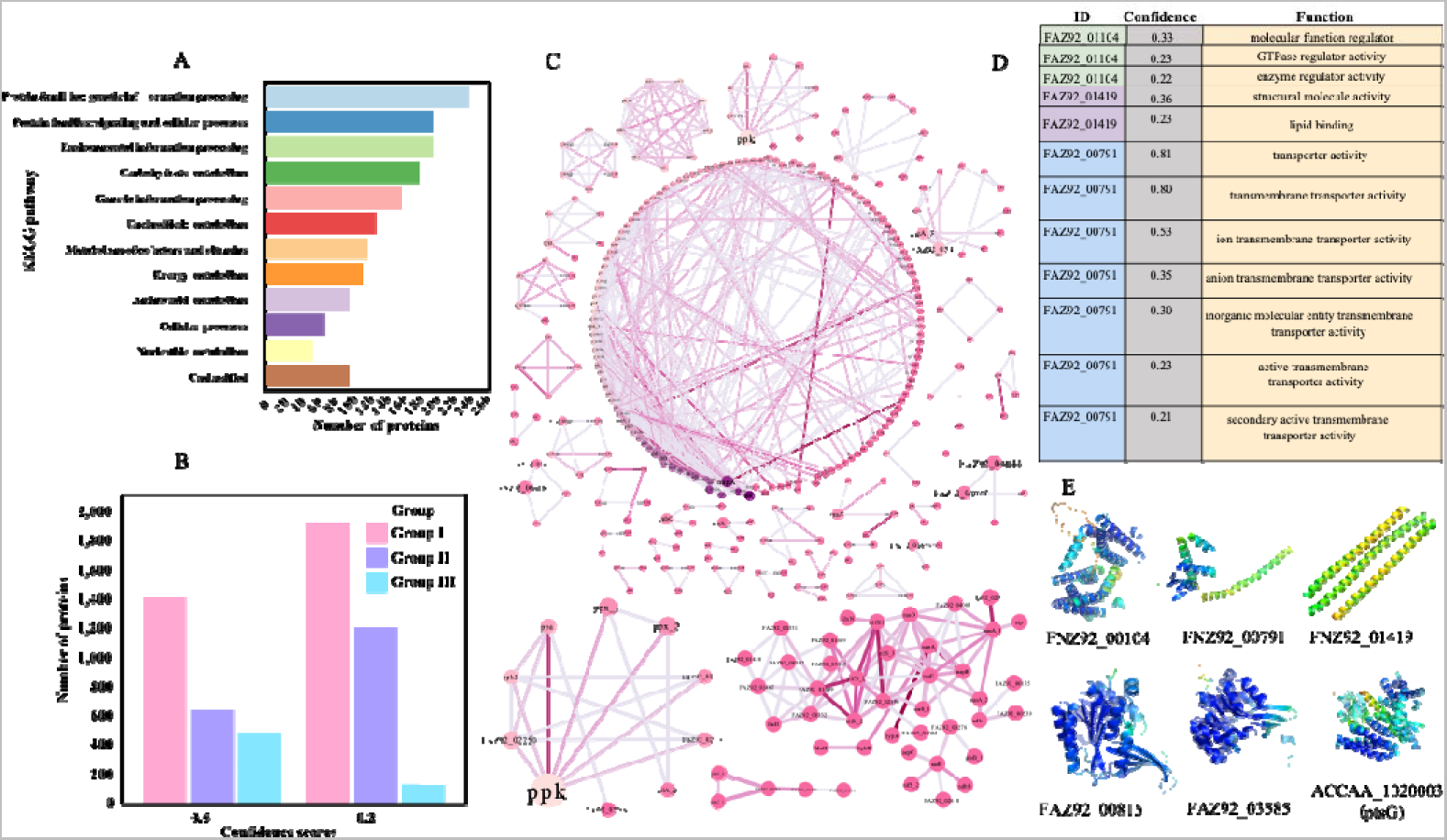
(A) The number of each functional category in SCELSE-1 based on KEGG. (B) Classifications of proteins in SCELSE-1 based on annotation results of KEGG and DeepFri. Proteins with successful or unsuccessful annotations were classified into three groups (Materials and Methods). (C) The protein-protein interactions in SCELSE-1 as predicted by STRING. (D) Functional predictions of hypothetical proteins in the PPI network (confidence >0.2). (E) The 3D structures of six carbon metabolism-related proteins.

### Evolution of core metabolic modules

A pangenome, constructed from 61 genomes encompassing 58 *Ca*. Accumulibacter, 1 *Ca*. Propionivibrio aalborgensis S1 (GAO) [50] and 2 *Ca*. Dechloromonas MAGs (PAOs) [49], yield 20,397 orthologous gene clusters (SI Spreadsheet 1). We employed Foldseek to predict structural orthologs in pan-structural proteomes (60,198) of *Ca*. Accumulibacter, uncovering 5,661 structural similar groups (structural orthologs). To better compare the genetic composition difference between PAOs and GAOs, we selected *Ca*. Accumulibacter similis SCELSE-1, and *Ca*. Propionivibrio aalborgensis S1 (a GAO) as representatives. Comparative analyses revealed that SCELSE-1 encoded 3,559 proteins (3,299 have predicted structures), of which 939 protein sequences and 1,301 protein structures were identified as similar to those in S1. These protein pairs were categorized into four groups based on their similarity/dissimilarity in functions, structures, and sequences: (I) non-orthologous protein sequences but with similar structures; (II) orthologous sequences that are structurally different; (III) orthologous sequences with similar structures; and, (IV) non-orthologous sequences with different structures. Grouped proteins were mapped to their associated metabolic pathways to understand their evolution (Figure 1D). Metabolic modules of different groups indicated distinct sources and evolutionary histories between PAO (represented by *Ca.* Accumulibacter similis SCELSE-1) and GAO (represented by *Ca*. Propionivibrio aalborgensis S1). In Group I, the key functions and structures of those proteins were preserved during evolution, although their encoding gene sequence may have changed rearrangements, insertions, deletions, etc. resulting in distinct gene sequences. Within Group I, proteins predominantly occurred in the ABC transporters and two-component system pathways, indicating that these proteins evolved into different genotypes, resulting in lower protein sequence similarities, whereas, similar functions were retained. Notably, protein pairs in Group II, characterized by having similar sequences but dissimilar structures, were consistently low, averaging at 1%. Group III exhibited the highest proportions in the TCA cycle (71%) and nitrogen metabolism (65%), implying the conservation of these metabolic pathways during evolution. Notably, central carbon metabolism pathways such as glycolysis/gluconeogenesis and the TCA cycle were highly conserved, and the vast majority of protein structures were highly conserved between *Ca*. Accumulibacter and *Ca*. Propionivibrio, regardless of the similarity of the encoding gene sequences (Figure 1E), representing conserved metabolic modules acquired from their common ancestor. In contrast, Group IV showed higher occurrences in quorum sensing and the pentose phosphate pathways, exceeding 30%. A complete pentose phosphate pathway was not encoded in SCELSE-1, whereas was preserved in S1. The disparity suggested that this pathway may have evolved independently after phylogenic differentiation, rather than a simple loss of genes during the evolution of *Ca*. Accumulibacter. The above results indicated that proteins occurred in a specific pathway appear to have evolved independently or as sub-units, rather than along with all other proteins within a specific pathway [44]. For example, the TCA cycle was typically treated as a whole in modern biological research. Whereas, proteins in this pathway distributed in different groups (Figure 1E). Even proteins that play the same role had undergone different evolutionary processes. For example, three acetyl-CoA synthetases in SCELSE-1 were classified into Group I, II, and III, respectively, suggesting their distinct evolution modes and histories.

### *Ca*. Accumulibacter phylogeny

Phylogenic analysis is a cornerstone in microbiological research. *Ca*. Accumulibacter are divided into 14 clades based on their *ppk*1 polymorphism [15]. Given the generally slower evolution rates of protein structures than gene sequences, we compared the phylogeny based on the PPK1 and PPK2structures. The encoding gene of the latter was laterally derived at the last common ancestor (LCA) of *Ca.* Accumulibacter, and was considered to play a key role in the emergence of the polyphosphate-accumulating trait of *Ca.* Accumulibacter [63]. Phylogenetic trees were built based on PPK1 and PPK2 structures, their encoding gene sequences, and the whole genome, to understand the evolutionary differences demonstrated by different markers (Figure 3). Both the whole-genome tree, the *ppk*1 and *ppk*2 gene trees exhibited robust resolutions, effectively grouping genomes from the same clade. Nevertheless, a notable incongruity emerged between the whole-genome and the *ppk*1/*ppk*2 trees, where Bin228 (clade II-I) displayed a closer evolutionary relationship to clade IIJ in the former, but a closer association with clade IIA in the latter. Remarkably, both PPK1 and PPK2 structure trees showed that Bin228 shared a closer evolutionary connection with clade IIA members, supporting a closer relationship between clades II-I and IIA.

**Figure 3.**
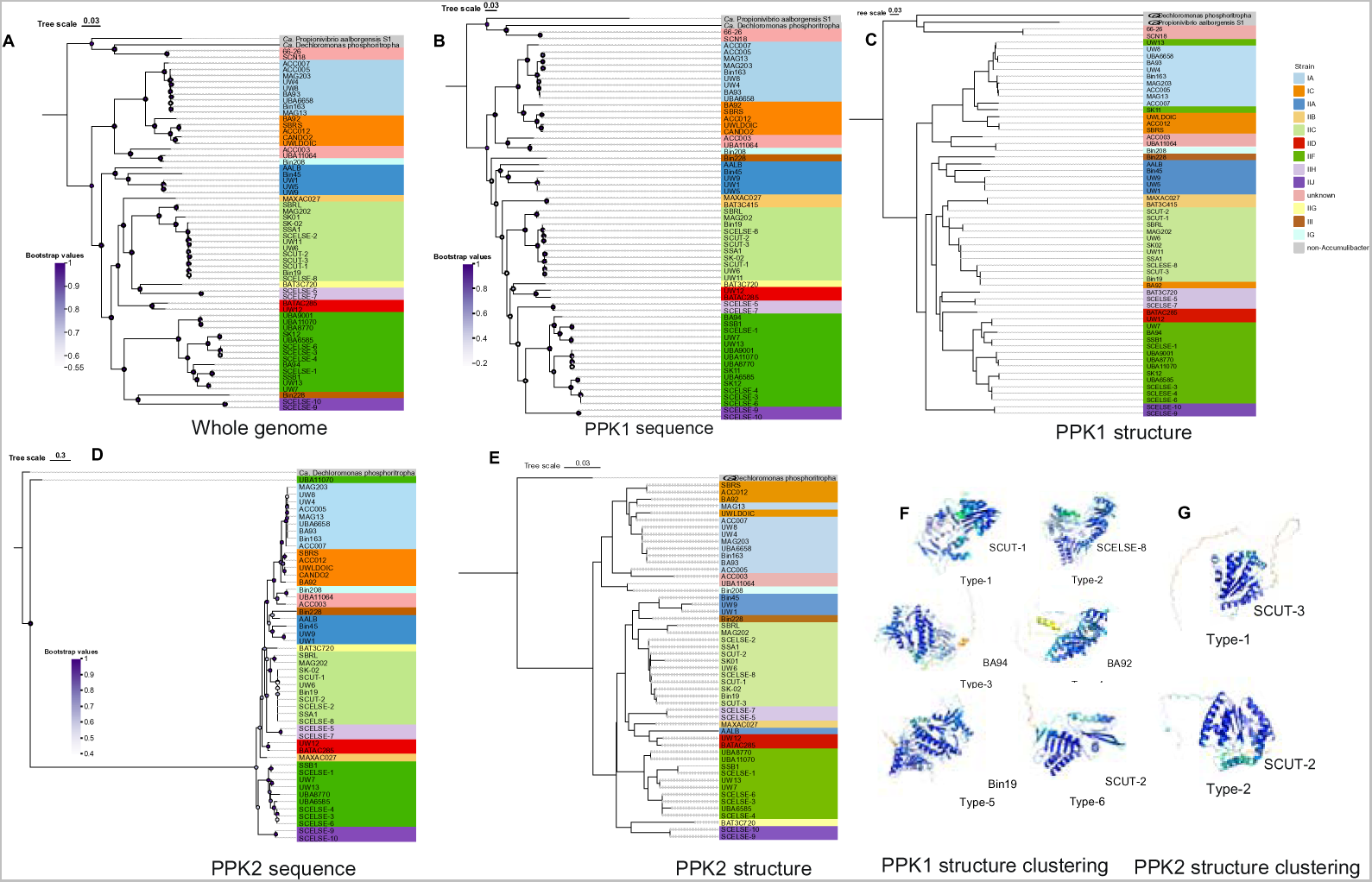
Phylogeny of *Ca*. Accumulibacter based on different markers: the whole genome (A), *ppk*1 sequence (B) and PPK1 structure (C), *ppk*2 sequence (D) and PPK2 structure (E). The phylogenetic trees of *Ca*. Accumulibacter were based on *ppk*2 and *ppk*1 gene sequences by using iqtree2. The phylogenetic trees based on PPK1 and PPK2 structures were constructed by using Foldtree. MAGs of *Ca*. Dechloromonas phosphoritropha (GCA_016722705.1) [49] and *Ca*. Propionivibrio aalborgensis S1 (GCA_900089945.1) were used as the outgroup [50]. (F) and (G) are the classification of PPK1 and PPK2 structures based on Foldseek.

Within the PPK1 structure tree, a majority of genomes from the same clade displayed cohesive clustering. Whereas, certain genomes showed unexpected associations. For instance, clade IIF member UW13 clustered into clade IA; clade IC member BA92 clustered into clade IIC, challenging *ppk*1-sequence base classification. Most members from the same clade also clustered together in the PPK2 structure tree, while a few members exhibit inconsistencies, e.g., clade IIA member AALB clustered into clade IID. To further elucidate the intricacies of PPK1 and PPK2 structures, Foldseek was employed for structure clustering. Although the *ppk*1 sequences uniformly clustered in one homologous gene cluster, structural similarity led to the division of PPK1 into six distinct types (Figure 3F). Notably, the Type-5 structural cluster encompassed the largest number of genomes, reaching 54 (61 in total), including *Ca*. Propionivibrio aalborgensis and *Ca*. Dechloromonas phosphoritropha. This clustering suggested an overall conserved structural evolution, as a majority of *Ca*. Accumulibacter genomes clustered together with their relatives within the Rhodocyclaceae family. However, within specific clades, high diversities in the PPK1 structures were evident. For instance, PPK1 in clade IIC members SCUT-1, SCUT-2, SCELSE-8, and Bin19 are allocated to different structure types. A intriguing question occurs: why do highly conserved sequences exhibit such pronounced structural differences? Previous studies by Kosloff et al. found several examples where two similar-sequence-encoding proteins had significantly different structures [64]. And functional significance that was related to structural differences between high-sequence-identity pairs. The conserved structure of PPK1 among *Ca*.

Accumulibacter PAO and *Ca*. Propionvibrio non-PAO and the nonconserved structures of PPK1 within *Ca.* Accumulibacter clade suggested that the *ppk*1 was not a key gene defining PAO and non-PAOs. In addition, PPK2 of *Ca*. Propionivibrio aalborgensis and *Ca*. Accumulibacter were not clustered based on sequences and structures, indicating that *ppk*2 is a differential gene between PAOs and GAOs. PPK2 structures were divided into two types based on Foldseek, which was more conservative than PPK1, indicating a relatively stable structure during evolution, and the essential role of its specific biological function (polyphosphate synthesis). These conserved sequences were also relatively conserved in structures, which may indicate that PPK2 better donated the evolution of *Ca*. Accumulibacter.

Interestingly, all trees showed that unknown clade members (66-26 and SCN18), which were previously classified as *Ca.* Accumulibacter based on 16S rRNA gene homology, were grouped closer to *Ca*. Dechloromonas, rather than other *Ca*. Accumulibacter members, demonstrating that 66-26, and SCN18 are not *Ca*. Accumulibacter members, highlighting that, for accurate and rational classification, the taxonomy based on 16S rRNA genes is to be combined with other biomarkers. Additionally, UBA11064 (clade identity unknown) and Bin208 were classified into a clade based on different marker genes, indicating that they belong to the same clade (i.e., clade IG members). The overall consistency with local differences in the results of structure- and sequence-base phylogenies indicated that the structure of the marker-gene-encoding proteins was both effective in displaying phylogeny and provided new perspectives for understanding the evolution of *Ca*. Accumulibacter, underscored the need for a multifaceted approach in microbial phylogenetics, and the potency of tertiary protein representation in decoding microbial phylogeny.

### Phosphorus metabolism

In the exploration of the distinct metabolisms of PAOs and GAOs, with a specific focus on P metabolism, intriguing findings have emerged. PPK2, a crucial enzyme for polyphosphate synthesis, has multiple (2-4) copies of encoding genes in *Ca.* Accumulibacter PAO and *Ca*. Propionvibrio GAO genomes. As a laterally derived gene at the *Ca.* Accumulibacter LCA, our prior research suggested the pivotal role of *ppk*2-1 in the emergence of the polyphosphate-accumulating feature of this group of bacteria [63]. Notably, the remaining 3 *ppk*2 copies in SCELSE-1 shared similar structures despite their dissimilar sequences to those in S1 (a GAO), suggesting functional similarities. The PPK2-1 belonged to Group IV (non-ortholog sequences and structures but similar functions) which was structurally and sequence different from other PPK2, further underscoring the uniqueness of PPK2-1 (Figure 1D) as a key marker distinguishing PAOs from GAOs.

The phosphate regulator (Pho) emerges as a key regulatory mechanism for maintaining and managing bacterial intracellular inorganic phosphate concentrations [65]. The signal transduction of Pho regulators requires PhoR, PhoB, and four components of the high-affinity phosphate ABC transporter (PstS, PstA, PstB, and PstC) [66, 67]. Our prior study identified multiple *pho*U homologs in pan-*Ca*. Accumulibacter genomes, which located near the *pit*, displayed increased transcription during an anaerobic-aerobic EBPR cycle [63]. By comparing their structures with those of existing proteins in the Alphafold database, although most matches were PhoU homologs, there is a putative *pit* accessory protein (Pap), the structure of which is largely similar to the PhoU homologs in *Ca.* Accumulibacter. PPI network analysis indicated interactions between the PhoU homologs and Pit, but not Pst, implying that the PhoU homologs in *Ca.* Accumulibacter were regulating Pit rather than Pst as in other bacteria (Figure 4A). Hdock-based predictions also showed binding sites between the Pit and the PhoU homolog (Figure 4C) with score and the average confidence score were −287.62 and 0.939, respectively. Whereas, those between the PhoU homolog and the PstS were −215.72 and 0.79, as low as those between PhoU and PstS (−209.35 and 0.76) (SI Figure S4 and Table S5), demonstrating the regulatory relationship between the PhoU homologs and the Pit instead of the Pst. To further testify the hypothesis, metatranscriptomic analysis was performed on enrichment cultures of *Ca.* Accumulibacter cognatus SCUT-2 and SCUT-3. Although the *pho*U homolog showed high transcription, the trend was not in line with *pst* (SI Figure S3), indicating that the PhoU homolog was not effectively regulating Pst. Within *Ca.* Accumulibacter cognatus SCUT-3, the *pho*U homolog-1 was located upstream *pit*-1; while *pho*U homolog -2 and -3 were located downstream *pit*-2 and 3, respectively (Figure 4B). The same phenomenon was widely observed in all *Ca*. Accumulibacter genomes. The transcriptional pattern of *pit*-1, inversely related to its PPI-associated *pho*U homolog, while those of *pit*-2 and *pit*-3 were positively correlated with their related *pho*U homologs, demonstrating the co-transcriptional patterns of the *pit*2- *pho*U homolog and *pit*3- *pho*U homolog systems but not the *pit*1-*pho*U homolog system. In *Streptomyces coelicolor* [68], a *pit* gene (namely *pit*H1) was located downstream *pap*, which formed a bicistronic with *pap* and was the main phosphate transporter under high extracellular phosphate concentration conditions. Another *pit* gene (named *pit*H2) was surrounded by genes that are not involved in phosphate uptake, and its transcription increased when phosphate was restricted. Given the similar transcriptional trends of *pit*-2 and *pit*-3 with their associated *pho*U homologs, despite their opposite arrangement patterns from *pit*H1 and *pap* in *Streptomyces coelicolor*, these *pho*U homologs were positively regulating their respective *pit*. Additionally, the transcription patterns of *pit*-1 in *Ca.* Accumulibacter was opposite the changes in the phosphate concentrations in the bioreactor (SI Figure S5), suggesting that this copy of *pit* is functionally similar to *pit*H2 in *Streptomyces coelicolor*. Overall, these observations prompted the assertion that PhoU homolog holds a regulatory relationship with Pit, rather than Pst, aligning with our earlier hypothesis that the polyphosphate-accumulating trait of *Ca.* Accumulibacter is a result of ineffective regulation of Pst by PhoU [63]. Overall, based on structural matching, PPI network, molecular docking and metatranscriptomic analysis, these PhoU homologs are identified as a Pit regulatory protein to having a similar function as Pap. These comprehensive approaches contributed to a novel and enhanced understanding of the intricate interplay of enzymes and regulatory mechanisms in the P removal process by *Ca*. Accumulibacter, offering valuable insights into PAO physiology.

**Figure 4.**
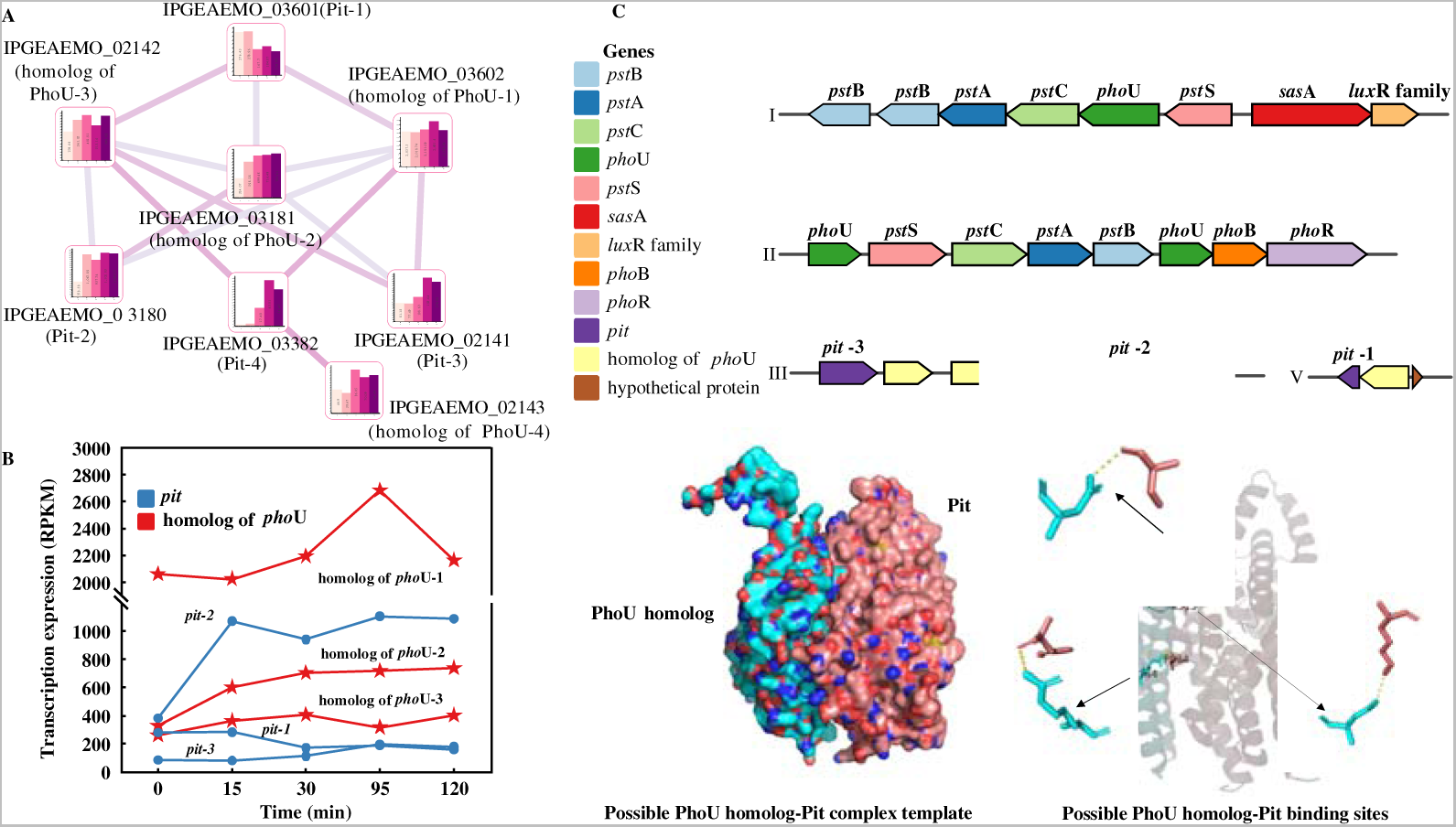
Protein-protein interactions in the P metabolic pathway in *Ca*. Accumulibacter cognatus SCUT-3. (A) The PPI network of *pit* and *pho*U homolog (now considered as *pap*) and (B) their transcriptional characteristics in SCUT-3. (C) The P metabolism-related gene cluster in SCUT-3. Genes are color-coded based on their functions. (D) Pap and Pit protein binding models and binding sites as predicted by Hdock.

### Other metabolism

The center carbon metabolism was considered to be highly conserved within the genus of *Ca*. Accumulibacter. Whereas, there was still a majority of inexplicableness. For example, glycolysis, an essential process for the generation of ATP and NADH for anaerobic carbon uptake and storage, could be achieved in two pathways (the Embden-Meyerhof-Parnas (EMP) pathway and the Entner-Doudor off (ED) pathway) [69, 70]. Notably, a careful examination of 47 high-quality *Ca*. Accumulibacte genomes suggested the lack of genes encoding the phosphofructokinase (PFK) in the EMP pathway in a great number of *Ca*. Accumulibacter genomes (36/47), albeit glycolysis is a highly conserved and essential metabolism of *Ca.* Accumulibacter. Via clustering analysis through Foldseek, two proteins FAZ92_00815 and FAZ92_03585 structurally clustered with the PFK protein in *Ca.* Accumulibacter aalborgensis AALB (Figure 2E). However, due to low sequence similarity, FAZ92_00815 was not clustered into the same orthogroup with *pfk* based on sequence-base annotation. Combined with sequence and structural clustering results, we found that the *pfk* gene was present in all *Ca.* Accumulibacter genomes (except MAXAC720), the previously observed absence of *pfk* in a majority of *Ca.* Accumulibacter genomes were a result of unsuccessful annotation by KEGG. Additionally, we found that only AALB and BAT3C720 (2/47) encoded the glucose-phosphoenolpyruvate phosphotransferase subunit G gene (*pts*G), which renders their unique ability to utilize glucose as a carbon source. By performing combined sequence and structure clustering analysis, no other genes were found to group with the *pts*G in AALB and BAT3C720, indicating that *pts*G was indeed absent in other *Ca*. Accumulibacter genomes rather than an annotation deficiency. The result further supported that glucose utilization was a specific metabolic capacity of a small number of *Ca*. Accumulibacter members (i.e., AALB and BAT3C720).

The confidence scores of a great number of nitrogen metabolism-related proteins, as suggested by DeepFri, were less than 0.2, indicating substantial differences from the DeepFRI training dataset, underscoring the existence of numerous unknowns in the nitrogen metabolism of *Ca.* Accumulibacter. Among proteins interacting with NirS or NapAB, FAZ92_01778 and FAZ92_07116 exhibited confidence scores higher than 0.2. These two proteins (interacting with NirS) displayed similar functional predictions, suggesting that they likely encode the same protein. The structural analogs using Foldsee identified their similarity to siroheme decarboxylase (a cofactor of NirS [71, 72]), which was consistent with the predicted function. Additionally, only clade IIF members SCELSE-3, 4 and 6, recovered from our high-temperature EBPR systems [46], were found to encode a complete denitrification pathway, as indicated by KEGG annotation. Three key proteins, which were commonly absent in other *Ca*. Accumulibacter genomes (i.e., NorC, NorB and NosZ), were predicted for structures by using Alphafold2 (with pLDDT > 90) and were put into the pan *Ca*. Accumulibacter structural genome for clustering. Based on their structures, a protein of 66-26 was clustered together with NosZ. This protein further clustered with a protein of SCN18 based on their encoding gene sequences. These results suggested that they may encode proteins with the same function as NosZ. Through structural annotation, these two genomes are suggested to encode a complete denitrification pathway, and the rest of *Ca*. Accumulibacter genomes lack the *nor*C gene, thus encode incomplete denitrification pathways.

By leveraging PPI networks and DeepFri analyses to elucidate the functions of hypothetical proteins in *Ca.* Accumulibacter, novel insights are obtained into key (P, carbon and nitrogen) pathways, highlighting the importance of structural analysis in refining genomic annotations and understanding microbial metabolism.

### Functional annotation reference database

Deep learning models, capable of capturing sequence spatial information and identifying functional residue, offer complementary information beyond traditional gene-based annotation reference. This suggested that a synergistic approach, combining deep learning with traditional sequence information, could expand the annotation coverage of the protein universe. Considering the ability to quickly and accurately annotate genomes was critical for the exploration of their encoding metabolic capacities, we have established a reference database based on Prokka (widely used for annotating *de novo* assemblies of prokaryotic genomes) to improve the resolution of annotation of *Ca.* Accumulibacter-related genomes [73]. We used 17,334 homolog gene clusters from the *Ca.* Accumulibacter pangenome which was constructed by curating 60 high-quality *Ca*. Accumulibacter genomes, where genes from each cluster were split into functions of known (1) or unknown (2). The clustering results were used to define the function of hypothetical proteins based on the hypothesis that genes from the same homologous gene cluster were considered to function identically. Moreover, the function of uncharacterized orthogroups was supplemented based on the functional annotation results of 60,189 protein structures with the highest confidence. The reference database (namely, “Accdate”) utilized the annotation information based on protein structures and gene sequences provided references for >200,000 *Ca*. Accumulibacter proteins. We used Accdata to annotate 28 *Ca*. Accumulibacter genomes and compared the results against the default database annotation, significantly improved the average annotation coverage to 83% from an average of <51% based on the default database (SI Figure 6). Our benchmarking of the Accdata showed that it was more accurate than either the structural or homologous gene annotation approach, minimizing unsuccessful annotations. In addition, the annotated results could be combined with PPI network analysis to provide more valuable metabolic insights such as those discussed in the P, carbon and nitrogen metabolism sections above. These results demonstrated that Accdata learned a meaningful reference for protein sequence annotation that (1) generalized to the dark matter of the sequence space and (2) used to predict and understand the properties of previously unknown gene-encoding proteins. A custom pipeline for generating different microbial annotation reference files was developed based on the above steps. The pipeline is readily applicable to any cultured and uncultured bacteria, establishing a link between microbial genes and protein structures, providing a more comprehensive annotated reference dataset for any bacteria of interest, facilitating the exploration of the gene dark matter in the bacteria domain.

## Credit authorship contribution statement

**Xiaojing Xie:** Conceptualization, Methodology, Software, Formal Analysis, Investigation, Data Curation, Writing - Original Draft, Visualization. **Xuhan Deng:** Data Curation, Resources, Visualization. **Liping Chen:** Data Curation, Resources, Visualization. **Jing Yuan:** Investigation, Resources, Data Curation. **Hang Chen:** Investigation, Resources, Data Curation. **Chaohai Wei:** Writing - Review & Editing, Supervision. **Xianghui Liu:** Investigation, Resources, Data Curation. **Chunhua Feng:** Supervision, Writing - Review & Editing. **Guanglei Qiu:** Conceptualization, Methodology, Investigation, Supervision, Writing - Review & Editing, Validation, Project Administration, Funding Acquisition.

## Declaration of Competing Interest

The authors declare that they have no known competing financial interests or personal relationships that could have appeared to influence the work reported in this paper.

## Acknowledgments

This research was supported by the National Natural Science Foundation of China (52270035 and 51808297), the Natural Science Foundation of Guangdong Province (2021A1515010494), the Guangzhou Key Research and Development Program (2023B03J1334), and the Pearl River Talent Recruitment Program (2019QN01L125).

## Data available

All data generated or analyzed during this study are included in this published article. Metagenomic raw reads and draft genomes were submitted to NCBI under the BioProject No. PRJNA807832, No. PRJNA994326 and No. PRJNA771771. Protein structure predictions are available at: https://github.com/XuhanDeng. Other data were documented in the Supplementary information.

## Code availability

DeepFRI v.0.0.1 for GO predictions (https://github.com/flatironinstitute/DeepFRI) and Alphafold2 for structure prediction (https://github.com/kalininalab/alphafold_non_docker). The structural clustering method is available at https://foldseek.com/, is implemented in Foldseek v.4.645b789 and is available as free and open-source software (GPLv3). The code for processing and analyzing and the pipeline used in this study has been uploaded to GitHub (https://github.com/XuhanDeng).

---

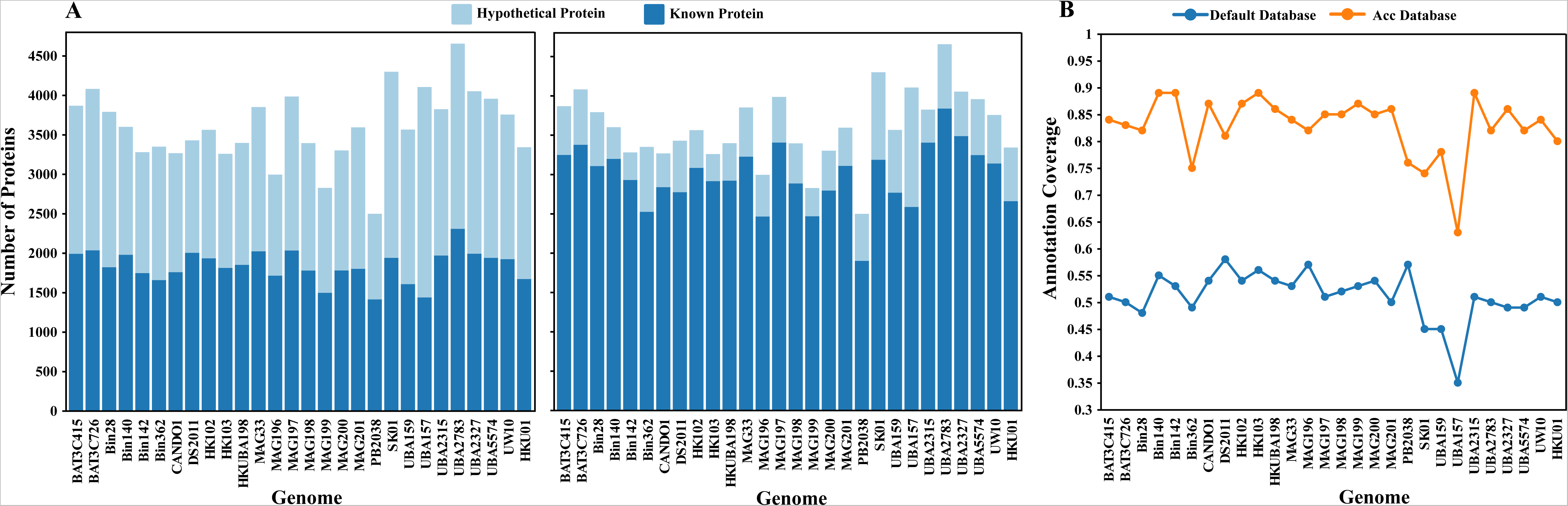

## Notes

https://github.com/XuhanDeng

### Competing Interest Statement

The authors have declared no competing interest.

