## Supplementary material for "From gene to structure: Unraveling genomic dark matter in *Ca*. Accumulibacter": Supplementary Material_EST.pdf

***Supplementary Methods***

**S1. *Ca.* Accumulibacter enrichment reactor operation**

The enrichment sequencing batch reactor (SBR) was inoculated in with activated sludge collected from a municipal wastewater treatment plant (WWTP) in Guangzhou, China. The SBR was operated with 6-h cycles, including a slow-feeding stage (60 min), an anaerobic stage (20 min), an aerobic stage (180 min), and a settling/decant stage (100 min). The SBR was fed with acetate as the sole carbon source, and a mineral salt medium with a phosphate stock solution to achieve a total organic carbon (TOC) to $PO_4^{3-}$-P ratio of 15:1 (C mol: P mol). In each cycle, two solutions, A and B, were introduced to both reactors at 2000 and 500 ml/cycle, respectively. Solution A is comprised of 4.3L of MilliQ™ water, 200 ml of sodium acetic trihydrates (113.3 g/L) and 500 ml of 10x nutrient stock solution (5L of MilliQ™ water, 10.2 g/L $NH_4Cl$, 12 g/L $MgSO_4 \cdot 7 H_2O$, 0.1 g/L peptone, 0.1 g/L yeast extract). Solution B is comprised of 25 L of MilliQ™ water, 13.75 ml of 200x phosphate stock solution (66 g/L $K_2HPO_4$, 51.5 g/L $KH_2PO_4$), 2ml of trace solution I (15 g/L of $FeCl_3 \cdot 6 H_2O$, 0.3 g/L of $CuSO_4 \cdot 5H_2O$, 0.6 g/L of $MnCl_2$, 1.2 g/L of $ZnSO_4$ and 1.5g/L of $CoCl_2$, acidified with 5ml of concentrated hydrochloric acid) and 1ml of trace solution II (1.5 g/L of $H_3BO_3$, 1.8 g/L of KI and 1.2 g/L of $Na_2MoO_4$). The hydraulic retention time (HRT) and sludge retention time (SRT) in the reactor were 12 h and 15 d, respectively. The pH was automatically controlled at 7.0-7.5 by using a M200 transmitter (Mettler-Toledo, Switzerland) connected to an acid/base (0.5 M HCl/NaOH) dosing system. The dissolved oxygen (DO) concentration was maintained at 0.8-1.2 mg/L during the aerobic phase by using the same transmitter connected to a solenoid valve in the aeration system. Temperature was controlled at 25°C, using a temperature controller connected to a thermostatic water bath. A programmable logic controller was linked to a SCADA interface for data visualization and storage.

**S2. Anaerobic-aerobic full cycle study**

**S2.1 Full-cycle study on the enrichment culture of *Ca.* Accumulibacter cognatus SCUT-2**

An anaerobic-aerobic full cycle study was performed on the enrichment culture of *Ca.* Accumulibacter cognatus SCUT-2 in the enrichment SBR at its original operation conditions with acetate as a carbon source as mentioned above. The full cycle contains an 80-min anaerobic stage (including a 60-min slow-feeding stage and a 20-min anaerobic stage afterwards) and a 180-min aerobic stage. Water and activated sludge samples was collected at 5, 15, 30, 45, 60, 80, 105, 120, 150, 180, 210 and 260 min for TOC, $PO_4^{3-}$-P, polyhydroxyalkanoates (PHA) and glycogen analyses. $PO_4^{3-}$-P concentrations were determined following the Standard Methods (APHA, 1999). TOC was measured using a TOC analyzer (Shimadzu, Japan). PHA analyses were performed according to Oehmen et al.[1], using a Trace Ultra Gas Chromatography (equipped with a DB-5MS column, 30 m ×0.25 mm, Agilent Technology, USA) coupled to a DSQ II mass spectrometer (Thermo Scientific, USA). Glycogen analyses were carried out according to the method described by Kristiansen et al.[2]. Lyophilized activated sludge was resuspended in 5 mL of 0.9 M HCl and digested at 100ºC for 5 h. Glucose equivalents in the supernatant were quantified using a high-performance liquid chromatography (E2695, Waters, US) equipped with a HyREZ XP column (Dionex, Thermo Fisher, Denmark). Activated sludge samples were collected just before the start of the full cycle (0 min), at 5 min (anaerobic phase), 15 min (anaerobic phase), 105 min (anaerobic phase), and 120 min (aerobic phase), snap-frozen in liquid $N_2$, and stored at -80°C before DNA/RNA extraction for metagenomic and metatranscriptomic analyses.

**S2.2 Full-cycle study on the enrichment culture of *Ca.* Accumulibacter cognatus SCUT-3**

Anaerobic-aerobic full- cycle study was performed in the SBR to investigate the dynamic gene transcriptions for *Ca.* Accumulibacter cognatus SCUT-3 under the reactor's original operating condition, except that 2.5L of synthetic wastewater was added at the beginning of the cycle as a pulse. Lactate was used to replace acetate in the influent wastewater to achieve the same TOC concentration. Other components remained unchanged. Filtered and mixed liquor samples were collected at regular time intervals until the end of each cycle for $PO_4^{3-}$-P, TOC, PHA, and glycogen analyses. For metagenomic and metatranscriptomic analysis, activated sludge samples were collected at 0 min (in the idle phase before cycle start, as control), 5 min (anaerobic phase), 15 min (anaerobic phase), 105 min (aerobic phase), and 120 min (aerobic phase), respectively. The samples were flash-frozen immediately in liquid nitrogen and stored at -80°C before DNA/RNA extraction.

**S3 Metagenomic analysis**

**S3.1 DNA extract and sequencing**

The first activated sludge samples (the ones at 0 min) collected in the above-mentioned full-cycle studies were used for metagenomic analyses. Genomic DNA were extracted using the OMEGA Soil DNA Kit (D5625-01) (Omega Bio-Tek, GA, USA), following the manufacturer's instructions, and stored at -80°C prior to further analyses. The quantity and quality of the extracted DNA were measured using a Qubit™ 4 Fluorometer (with WiFi: Q33238; Qubit™ Assay Tubes: Q32856; Qubit™ 1X dsDNA HS Assay Kit: Q33231) and 1% agarose gel electrophoresis, respectively. Genomic library was constructed following the Illumina TruSeq DNA Sample Preparation Guide. Illumina NovaSeq sequencing run at a read-length of 150 bp paired-end. Low-quality reads and adapter sequences were removed using fastp 0.20.1 [3]. metaSPAdes v.3.13.0 was used to assemble high-quality reads to contigs [4]. The contigs were binned into metagenome-assembled genomes (MAGs) using MetaBAT2 v1.7 and CONCOCT v1.1 [5, 6]. CheckM v1.0.18 was used to evaluate the completeness and contamination of MAGs[7]. The taxonomic classifications of obtained MAGs were performed using the Genome Taxonomy Database Toolkit (GTDB-Tk v. 2.3.0) [8]. MAGs with completeness below 95% and contamination over 6% were discarded. BBMap v38.96 was used to calculate the relative abundance of each *Ca*. Accumulibacter MAGs [9]. All MAGs were annotated using RAST annotation server [10]. Except for quality control by fastp, annotation by RAST and relative abundance calculation by BBMap and taxonomic classifications for obtained MAGs by the GTDB-TK, all analyses, including metaSPAdes, MetaBAT2, COCOCT, CheckM, were performed on the KBase platform [11]. Raw reads and MAGs have been submitted to the National Center for Biotechnology Information (NCBI) and are accessible under BioProject No. PRJNA807832, No. PRJNA 994326, and No. PRJNA771771.

**S3.2 Metatranscriptomic analysis**

Activated sludge samples collected during the above-mentioned full-cycle studies were used for metatranscriptomic analyses. Total RNA was extracted using the RNA PowerSoil® Total RNA

Isolation Kit (Omega Bio-Tek, GA, USA). The quality and quantity of the extracted RNA were measured using 1.5% agarose gel electrophoresis and UV spectrophotometer, respectively. cDNA library was constructed using a TruSeq Standard mRNA LT Sample Prep Kit (Illumina, CA, USA). The quality and quantity of the library were measured using Agilent Bioanalyzer and Promega QuantiFluor, respectively. 10 ng RNA was loaded for each sample. Illumina NovaSeq sequencing was performed at a read-length of 150 bp paired-end. fastp [3] and SortMeRNA [12] were used to remove adaptation sequences and ribosomal ribonucleic acids (rRNAs). Filtered reads were mapped to the corresponding *Ca.* Accumulibacter MAGs (i.e., SCUT-2 or SCUT-3) using BBMap version 38.96 (9) and were normalized to transcript per million (TPM) and/or Reads Per Kb-CDSs Per Million Mapped Reads (RPKM). Raw reads were submitted to NCBI under the BioProject No. PRJNA771771.

***Supplementary Tables***

**Table S1** The EBPR performances and activated sludge characterizations of the enrichment cultures at the operational cycle of sludge sampling for metagenomic analyses

| Enrichment culture | Operational parameters | Measurements at the operational cycle |
|---|---|---|
| **SCUT-2** | Influent $PO_4^{3-}$-P (mg/L) | 14±3 |
| | Influent TOC (mg/L) | 58±3 |
| | MLSS (g/L) | 3.0 |
| | MLVSS (g/L) | 2.25 |
| | Anaerobic P release (mg P/g SS) | 36.67 |
| | Effluent $PO_4^{3-}$-P (mg/L) | 0.2~0.5 |
| **SCUT-3** | Influent $PO_4^{3-}$-P (mg/L) | 14±3 |
| | Influent TOC (mg/L) | 58±3 |
| | MLSS (g/L) | 3.0 |
| | MLVSS (g/L) | 2.25 |
| | Anaerobic P release (mg P/g SS) | 31.8 |
| | Effluent $PO_4^{3-}$-P (mg/L) | 0.2~0.5 |

Abbreviations: TOC, total organic carbon; MLSS, mixed suspended solid; VSS, volatile suspended solid

**Table S2** Information of MAGs used in this study. Genomic information includes clade classification, assembly accession, completeness and contamination.

| Strain | Species name | Clade | Accession | Completeness | contamination | Structure[1] |
|---|---|---|---|---|---|---|
| BA-93 | *Ca*. Accumulibacter regalis | IA | GCA_000585075.1 | 100 | 0.89 | + |
| UW8 | *Ca*. Accumulibacter regalis | IA | GCA_017302345.1 | 100 | 2.47 | - |
| UBA6658 | *Ca*. Accumulibacter regalis | IA | GCA_002455435.1 | 99.85 | 0.22 | - |
| ACC007 | *Ca*. Accumulibacter regalis | IA | GCA_907163205.1 | 100 | 4.89 | - |
| ACC005 | *Ca*. Accumulibacter regalis | IA | GCA_907163215.1 | 98.29 | 5.23 | - |
| MAG-203 | *Ca*. Accumulibacter regalis | IA | GCA_016790605.1 | 99.44 | 1.5 | - |
| SBRS | *Ca*. Accumulibacter delftensis | IC | GCA_012939955.1 | 97.11 | 4.96 | - |
| ACC012 | *Ca*. Accumulibacter delftensis | IC | GCA_907163155.1 | 99.96 | 4.6 | - |
| UWLDOIC | *Ca*. Accumulibacter meliphilus | IC | GCA_003332265.1 | 99.98 | 3.67 | - |
| UW1 | *Ca*. Accumulibacter phosphatis | IIA | GCA_000024165.1 | 99.99 | 0.41 | + |
| AALB | *Ca*. Accumulibacter aalborgensis | IIA | GCA_900089955.1 | 100 | 0.2 | + |
| UW9 | *Ca*. Accumulibacter phosphatis | IIA | GCA_017302455.1 | 99.98 | 3.67 | - |
| UW5 | *Ca*. Accumulibacter phosphatis | IIA | GCA_017592745.1 | 97.04 | 7.78 | - |
| MAXAC027 | *Ca*. Accumulibacter propinquus | IIB | GCA_016712935.1 | 99.99 | 3.99 | - |
| BAT3C415 | *Ca*. Accumulibacter propinquus | IIB | GCA_016714935.1 | 91.13 | 1.16 | - |
| SSA1 | *Ca*. Accumulibacter cognatus | IIC | GCA_013414765.1 | 100 | 0.08 | + |
| UW6 | *Ca*. Accumulibacter cognatus | IIC | GCA_017592725.1 | 99.99 | 3.71 | + |
| Bin19 | *Ca*. Accumulibacter cognatus | IIC | GCA_005889575.1 | 99.97 | 1.5 | - |
| SK-02 | *Ca*. Accumulibacter cognatus | IIC | GCA_000584975.2 | 99.99 | 0.03 | - |
| SCUT-1 | *Ca*. Accumulibacter cognatus | IIC | GCA_020709745.1 | 100 | 1.11 | - |
| SBRL | *Ca*. Accumulibacter contiguus | IIC | GCA_012940005.1 | 98.59 | 1.45 | - |
| SCELSE-2 | *Ca*. Accumulibacter cognatus | IIC | GCA_023806705.1 | 95.6 | 3.75 | - |

| | | | | | | |
|---|---|---|---|---|---|---|
| SCELSE-8 | *Ca.* Accumulibacter cognatus | IIC | GCA_023806525.1 | 91.84 | 0.5 | - |
| SCUT-3 | *Ca.* Accumulibacter cognatus | IIC | PRJNA993120 | 97.14 | 1.85 | - |
| SCUT-2 | *Ca.* Accumulibacter cognatus | IIC | GCA_024380085.1 | 100 | 1.11 | - |
| UW12 | *Ca.* Accumulibacter necessarius | IID | GCA_017302435.1 | 99.99 | 0.6 | - |
| BATAC285 | *Ca.* Accumulibacter proximus | IID | GCA_016709675.1 | 99.86 | 7.89 | - |
| SSB1 | *Ca.* Accumulibacter similis | IIF | GCA_013347225.1 | 100 | 0.23 | + |
| UW7 | *Ca.* Accumulibacter conexus | IIF | GCA_017592775.1 | 100 | 2.78 | - |
| UW13 | *Ca.* Accumulibacter conexus | IIF | GCA_017302415.1 | 100 | 7.14 | - |
| UBA6585 | *Ca.* Accumulibacter | IIF | GCA_002433845.1 | 98.98 | 0.81 | - |
| SCELSE-1 | *Ca.* Accumulibacter similis | IIF | GCA_005524045.1 | 97.11 | 4.96 | + |
| SCELSE-3 | *Ca.* Accumulibacter | IIF | GCA_023806635.1 | 99.99 | 0.62 | - |
| SCELSE-4 | *Ca.* Accumulibacter | IIF | GCA_023806545.1 | 98.18 | 0.52 | - |
| SCELSE-6 | *Ca.* Accumulibacter | IIF | GCA_023806505.1 | 93.64 | 0.53 | - |
| SCELSE-5 | *Ca.* Accumulibacter tropicus | IIH | GCA_023806585.1 | 99.05 | 0.63 | - |
| SCELSE-7 | *Ca.* Accumulibacter tropicus | IIH | GCA_023806565.1 | 99.85 | 1.01 | - |
| SCELSE-9 | *Ca.* Accumulibacter torridus | IIJ | GCA_023806595.1 | 99.99 | 0.92 | - |
| SCELSE-10 | *Ca.* Accumulibacter torridus | IIJ | GCA_023806625.1 | 100 | 1.09 | - |
| 66-26 | *Ca.* Accumulibacter sp | unknown | GCA_001897745.1 | 100 | 0.93 | + |
| SCN18 | *Ca.* Accumulibacter | unknown | GCA_017307795.1 | 100 | 1.11 | - |
| ACC003 | *Ca.* Accumulibacter | unknown | GCA_907163085.1 | 100 | 0.71 | - |
| BAT3C720 | *Ca.* Accumulibacter affinis | IIG | GCA_016713625.1 | 95.71 | 2.87 | - |
| Bin163 | *Ca.* Accumulibacter | IA | GCA_937891375.1 | 100 | 0.59 | - |
| Bin45 | *Ca.* Accumulibacter | IIA | GCF_029255285.1 | 100 | 0.69 | - |
| Bin228 | *Ca.* Accumulibacter | III | GCA_029255125.1 | 99.96 | 0.78 | - |
| MAG13 | *Ca.* Accumulibacter | IA | GCA_027492785.1 | 100 | 1.35 | - |
| Bin208 | *Ca.* Accumulibacter | IG | GCA_029255145.1 | 99.99 | 0.14 | - |
| UW11 | *Ca.* Accumulibacter | IIC | GCA_017302385.1 | 98.36 | 0.17 | - |

| | | | | | | |
|---|---|---|---|---|---|---|
| MAG202 | | IIC | GCA_016790665.1 | 99.05 | 0.63 | - |
| UW4 | *Ca*. Accumulibacter regalis | IA | GCA_017592785.1 | 95.6 | 3.75 | - |
| UBA11070 | *Ca*. Accumulibacter clone | IIF | GCA_003535635.1 | 68.34 | 0.08 | + |
| UBA11064 | *Ca*. Accumulibacter clone | unknown | GCA_003538495.1 | 84.26 | 8.03 | + |
| UBA8770 | *Ca*. Accumulibacter clone | IIF | GCA_003487685.1 | 74.79 | 0.35 | + |
| SK11 | *Ca*. Accumulibacter | IIF | GCA_000584995.1 | 79.91 | 0.39 | + |
| BA92 | *Ca*. Accumulibacter appositus | IC | GCF_000585055.1 | 93.93 | 0.74 | + |
| BA94 | *Ca*. Accumulibacter | IIF | GCA_000585095.1 | 74.06 | 7.56 | + |
| UBA9001 | *Ca*. Accumulibacter clone | IIF | GCA_003542235.1 | 58.88 | 0.2 | + |
| SK12 | *Ca*. Accumulibacter adjunctus | IIF | GCA_000585015.1 | 93.42 | 0.83 | + |
| CANDO2 | *Ca*. Accumulibacter | IC | GCA_009467885.1 | 86.63 | 1.44 | + |

[1] indicates the presence/absence of 3D structures of proteins encoded in each MAGs in Alphafold database. "+" indicates presence. "-" represents absence. The items color-coded in red are those recovered from our enrichment reactors.

**Table S3** The corresponding TAX ID of *Ca*. Accumulibacter in the Alphafold database.

| Number | Package name | Tax ID |
|---|---|---|
| 1 | proteome-tax_id-522306-0_v4.tar | 522306 |
| 2 | proteome-tax_id-2053492-0_v4.tar | 2053492 |
| 3 | proteome-tax_id-2053492-1_v4.tar | 2053492 |
| 4 | proteome-tax_id-2053492-2_v4.tar | 2053492 |
| 5 | proteome-tax_id-1860102-0_v4.tar | 1860102 |
| 6 | proteome-tax_id-327160-0_v4.tar | 327160 |
| 7 | proteome-tax_id-327160-1_v4.tar | 327160 |
| 8 | proteome-tax_id-1454003-0_v4.tar | 1454003 |
| 9 | proteome-tax_id-1454001-0_v4.tar | 1454001 |
| 10 | proteome-tax_id-1895689-0_v4.tar | 1895689 |
| 11 | proteome-tax_id-1454000-0_v4.tar | 1454000 |

Genome were obtained from the Alphafold database by running gsutil -m cp gs://public-datasets-deepmind-alphafold-v4/proteomes/proteome-tax_id-[TAX ID]-*_v4.tar

**Table S4** The structural clustering results of PPK1 and PPK2 using FoldSeek [13]. The PPK1 structures are divided into six types, and the PPK2 structures were divided into two types.

| Genome | Clade | PPK1 | PPK2 |
|---|---|---|---|
| SCUT-1 | IIC | Type1 | Type 2 |
| SCELSE-8 | IIC | Type2 | Type 2 |
| BA94 | IIF | Type3 | - |
| BA92 | IC | Type4 | Type 1 |
| *Ca*. Dechloromonas phosphoritropha | non-Accumulibacter | Type5 | Type 2 |
| Bin19 | IIC | Type5 | Type 1 |
| BA-93 | IA | Type6 | Type 1 |
| UW7 | IIF | Type5 | Type 2 |
| UW12 | IID | Type5 | Type 2 |
| ACC005 | IA | Type5 | Type 2 |
| ACC007 | IA | Type5 | Type 2 |
| MAXAC027 | IIB | Type5 | Type 2 |
| SK-02 | IIC | Type5 | Type 1 |
| SBRS | IC | Type5 | Type 1 |
| UW13 | IIF | Type5 | Type 2 |
| UW1 | IIA | Type5 | Type 1 |
| SBRL | IIC | Type5 | Type 1 |
| BATAC285 | IID | Type5 | Type 2 |
| UBA6585 | IIF | Type5 | Type 1 |
| BAT3C415 | IIB | Type5 | - |
| MAG-203 | IA | Type5 | Type 2 |
| UWLDOIC | IC | Type5 | Type 1 |
| SCELSE-1 | IIF | Type5 | Type 1 |
| SCELSE-3 | IIF | Type5 | Type 2 |
| SCELSE-4 | IIF | Type5 | Type 2 |
| UW8 | IA | Type5 | Type 2 |
| SCELSE-5 | IIH | Type5 | Type 2 |
| SCELSE-6 | IIF | Type5 | Type 2 |
| SCELSE-7 | IIH | Type5 | Type 2 |
| SCELSE-9 | IIJ | Type5 | Type 2 |
| SCELSE-10 | IIJ | Type5 | Type 2 |
| SCUT-3 | IIC | Type5 | Type 1 |

| | | | |
|---|---|---|---|
| ACC012 | IC | Type5 | Type 2 |
| AALB | IIA | Type5 | Type 1 |
| BAT3C720 | IIG | Type5 | Type 1 |
| Bin45 | IIA | Type5 | Type 2 |
| Bin228 | III | Type5 | Type 2 |
| MAG13 | IA | Type5 | Type 2 |
| UW11 | IIC | Type5 | - |
| MAG202 | IIC | Type5 | Type 2 |
| UW9 | IIA | Type5 | Type 1 |
| UW4 | IA | Type5 | Type 2 |
| UBA11070 | IIF | Type5 | Type 2 |
| UBA8770 | IIF | Type5 | Type 2 |
| UBA9001 | IIF | Type5 | - |
| SK12 | IIF | Type5 | - |
| UBA6658 | IA | Type5 | Type 1 |
| Bin163 | IA | Type5 | Type 1 |
| SSA1 | IIC | Type5 | Type 2 |
| SSB1 | IIF | Type5 | Type 2 |
| UW5 | IIA | Type5 | - |
| SCN18 | unknown | Type5 | - |
| 66-26 | unknown | Type5 | - |
| UBA11064 | unknown | Type5 | Type 2 |
| *Ca*. Propionivibrio aalborgensis | non-Accumulibacter | Type5 | - |
| ACC003 | unknown | Type5 | Type 1 |
| Bin208 | IG | Type5 | Type 1 |
| SK11 | IIF | Type5 | - |
| SCUT-2 | IIC | Type6 | Type 2 |
| UW6 | IIC | Type2 | Type2 |

**Table S5** The complex template information of different proteins **(**summary of the top 10 models). The docking scores are calculated by knowledge-based iterative scoring function ITScorePP or ITScorePR. A more negative docking score means a more possible binding model. When the confidence score was above 0.7, the two molecules would be very likely to bind; When the confidence score was between 0.5 and 0.7, the two molecules would be possible to bind; when the confidence score is below 0.5, the two molecules would be unlikely to bind.  The ligand RMSDs are calculated by comparing the ligands in the docking models with the input or modeled structures.

| **PhoU homolog-Pit** | | | | | | | | | | | |
|---|---|---|---|---|---|---|---|---|---|---|---|
| Rank | **1** | **2** | **3** | **4** | **5** | **6** | **7** | **8** | **9** | **10** | Average |
| Docking Score | -310.8 | -292.4 | -290.3 | -290 | -287.5 | -284.5 | -283.3 | -279.8 | -278.8 | -278.8 | -287.625 |
| Confidence Score | 0.9614 | 0.9452 | 0.943 | 0.9426 | 0.9399 | 0.9365 | 0.935 | 0.9307 | 0.9293 | 0.9293 | 0.93929 |
| Ligand rmsd (Å) | 52.48 | 39.52 | 35.07 | 54.3 | 46.13 | 48.12 | 48.97 | 41.96 | 45.28 | 44.12 | 45.595 |
| **PhoU homolog-PstS** | | | | | | | | | | | |
| Rank | 1 | 2 | 3 | 4 | 5 | 6 | 7 | 8 | 9 | 10 | Average |
| Docking Score | -238.1 | -223.1 | -222.1 | -217.7 | -215 | -211.1 | -210 | -209.3 | -206.3 | -204.6 | -215.727 |
| Confidence Score | 0.8534 | 0.8119 | 0.8089 | 0.7948 | 0.7859 | 0.7723 | 0.7684 | 0.7661 | 0.7552 | 0.7486 | 0.78655 |
| Ligand rmsd (Å) | 46.53 | 49.65 | 47.72 | 45.47 | 42.87 | 40.31 | 36.72 | 52.26 | 51.02 | 48.68 | 46.123 |
| **PhoU-PstS** | | | | | | | | | | | |
| Rank | 1 | 2 | 3 | 4 | 5 | 6 | 7 | 8 | 9 | 10 | Average |
| Docking Score | -227.5 | -225.1 | -217.3 | -210.6 | -205.5 | -204 | -201.9 | -201.5 | -201.1 | -199.2 | -209.353 |
| Confidence Score | 0.8249 | 0.8178 | 0.7933 | 0.7705 | 0.7521 | 0.7466 | 0.7384 | 0.7368 | 0.7352 | 0.7277 | 0.76433 |
| Ligand rmsd (Å) | 43.13 | 33.03 | 49.73 | 42.6 | 42.84 | 49.27 | 44.21 | 53.3 | 56.35 | 43.15 | 45.761 |

### *Supplementary Figures*

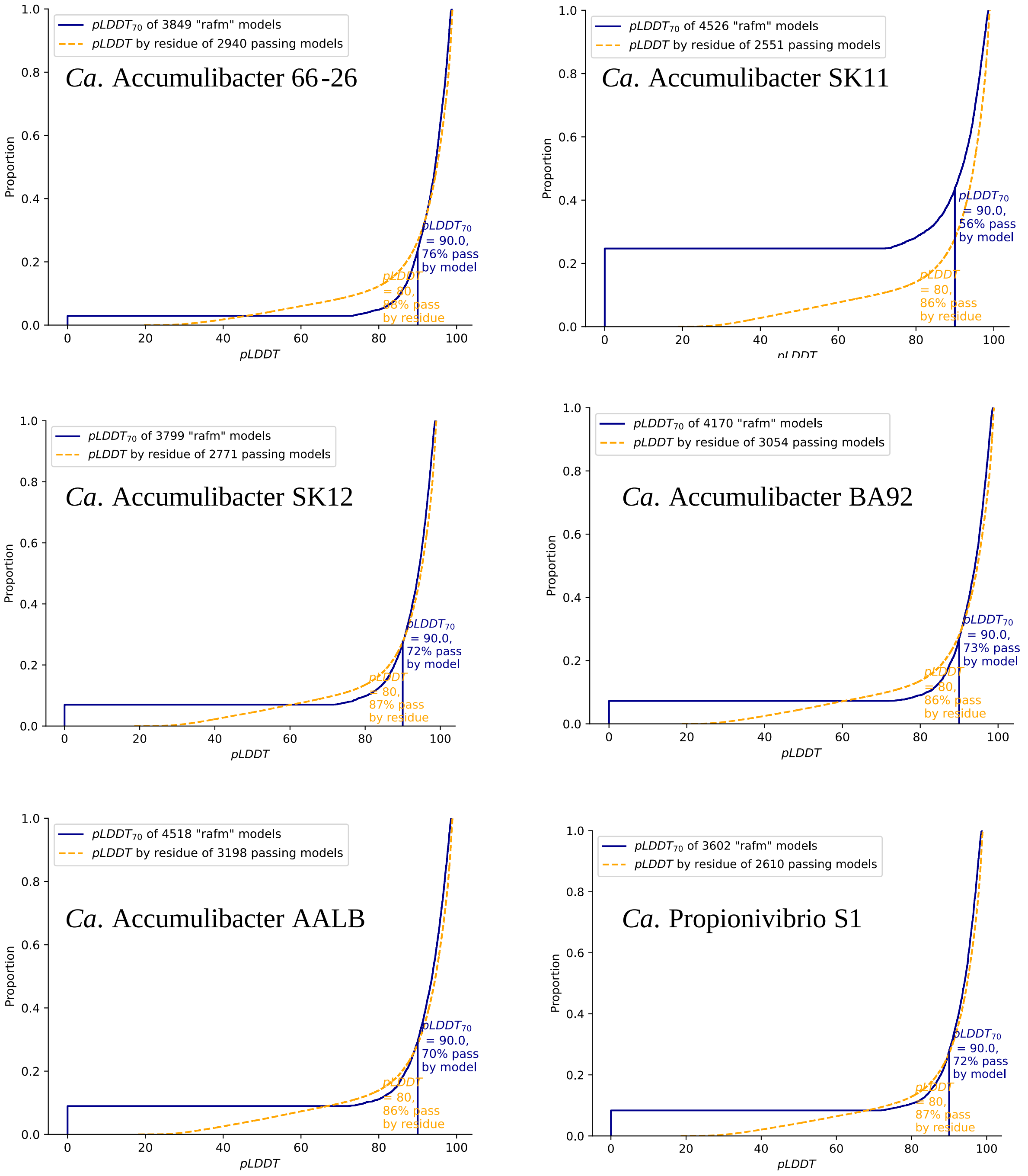

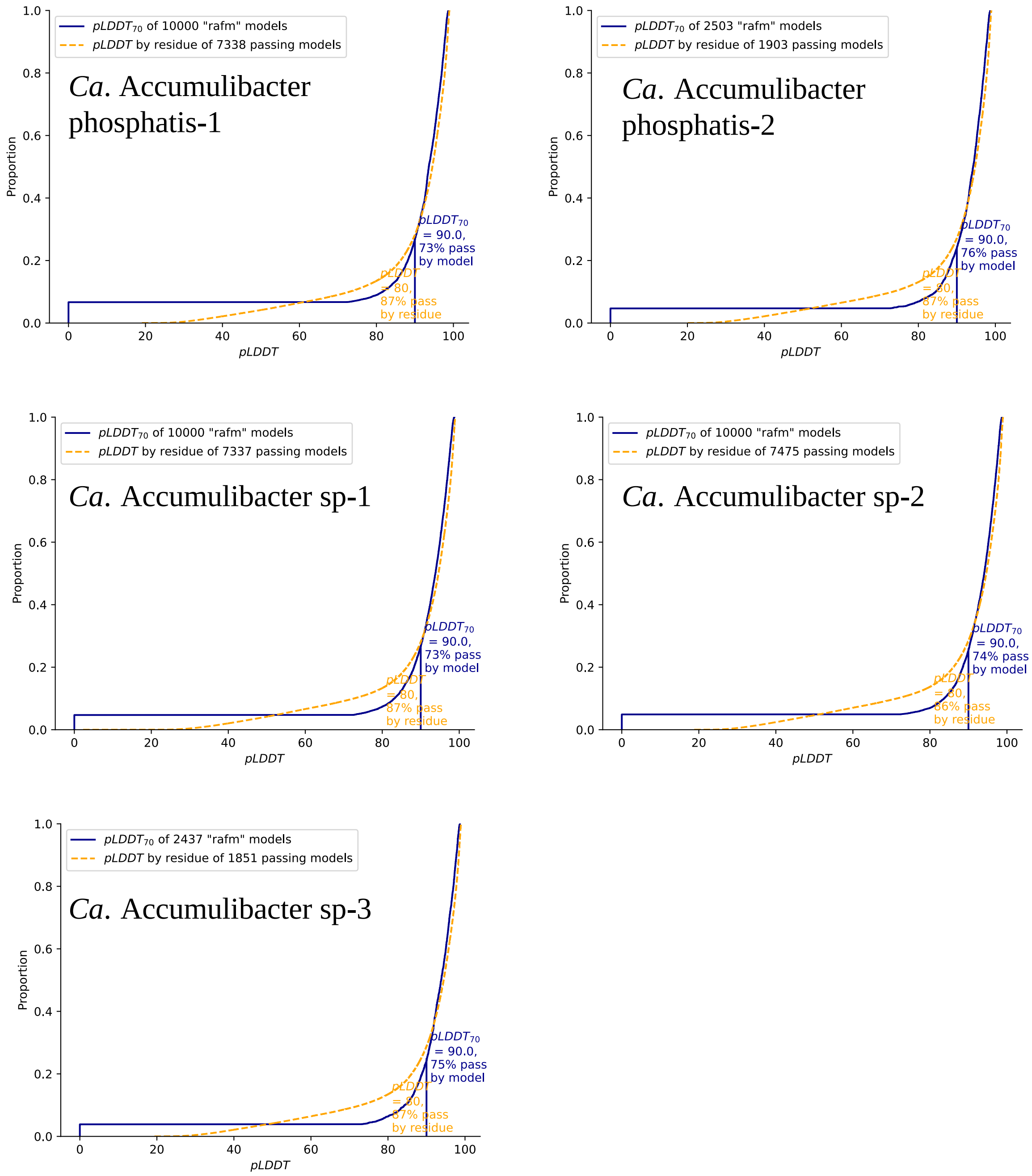


**Figure S1** Quality assessment and clustering of protein structures. The pLDDT confidence scores of pre-models which were obtained from the Alphafold database. The distributions of the bounded pLDDT and residues in models are shown in each sub-pannels.

A

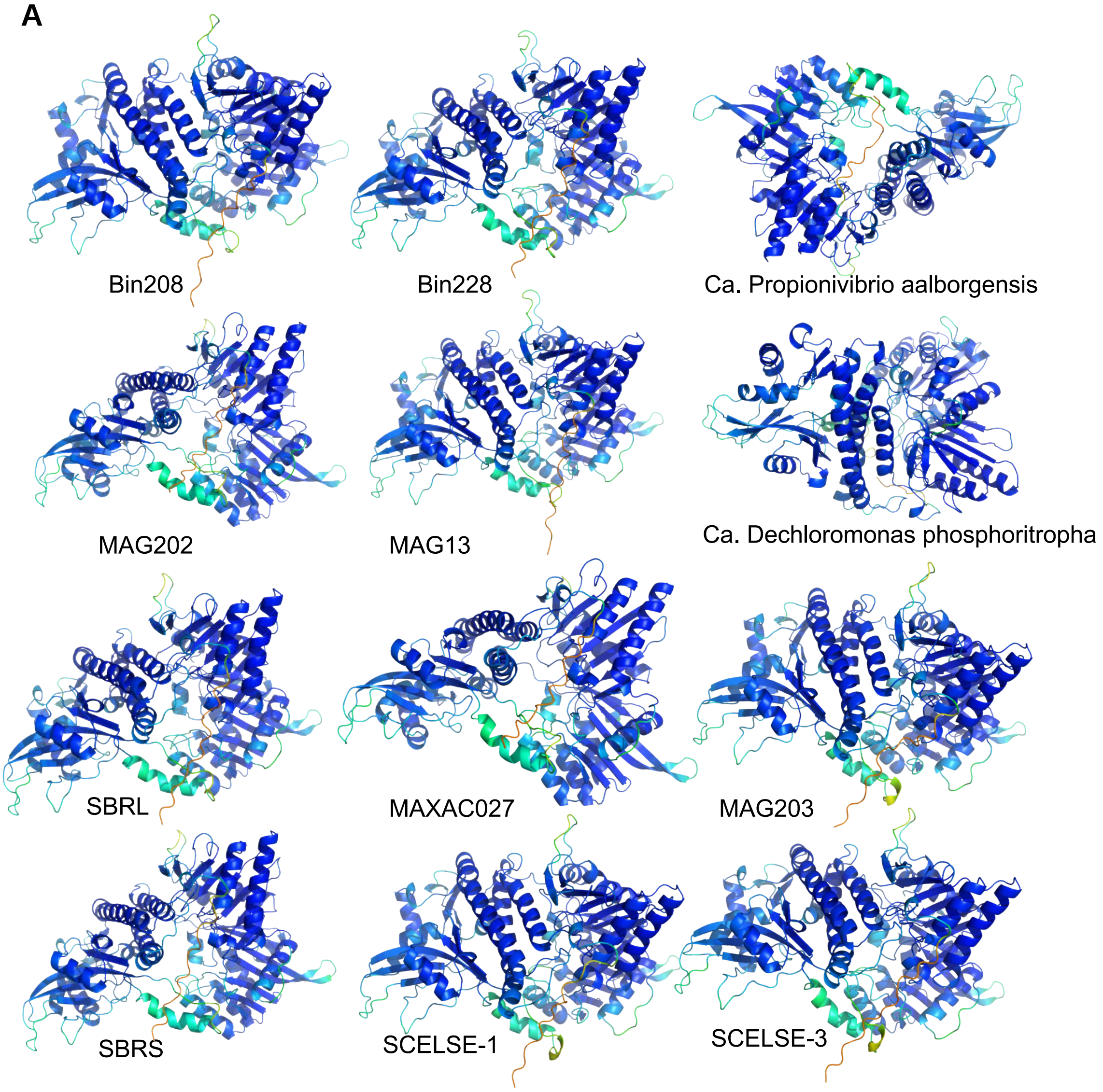
Bin208
Bin228
Ca. Propionivibrio aalborgensis
MAG202
MAG13
Ca. Dechloromonas phosphoritropha
SBRL
MAXAC027
MAG203
SBRS
SCELSE-1
SCELSE-3

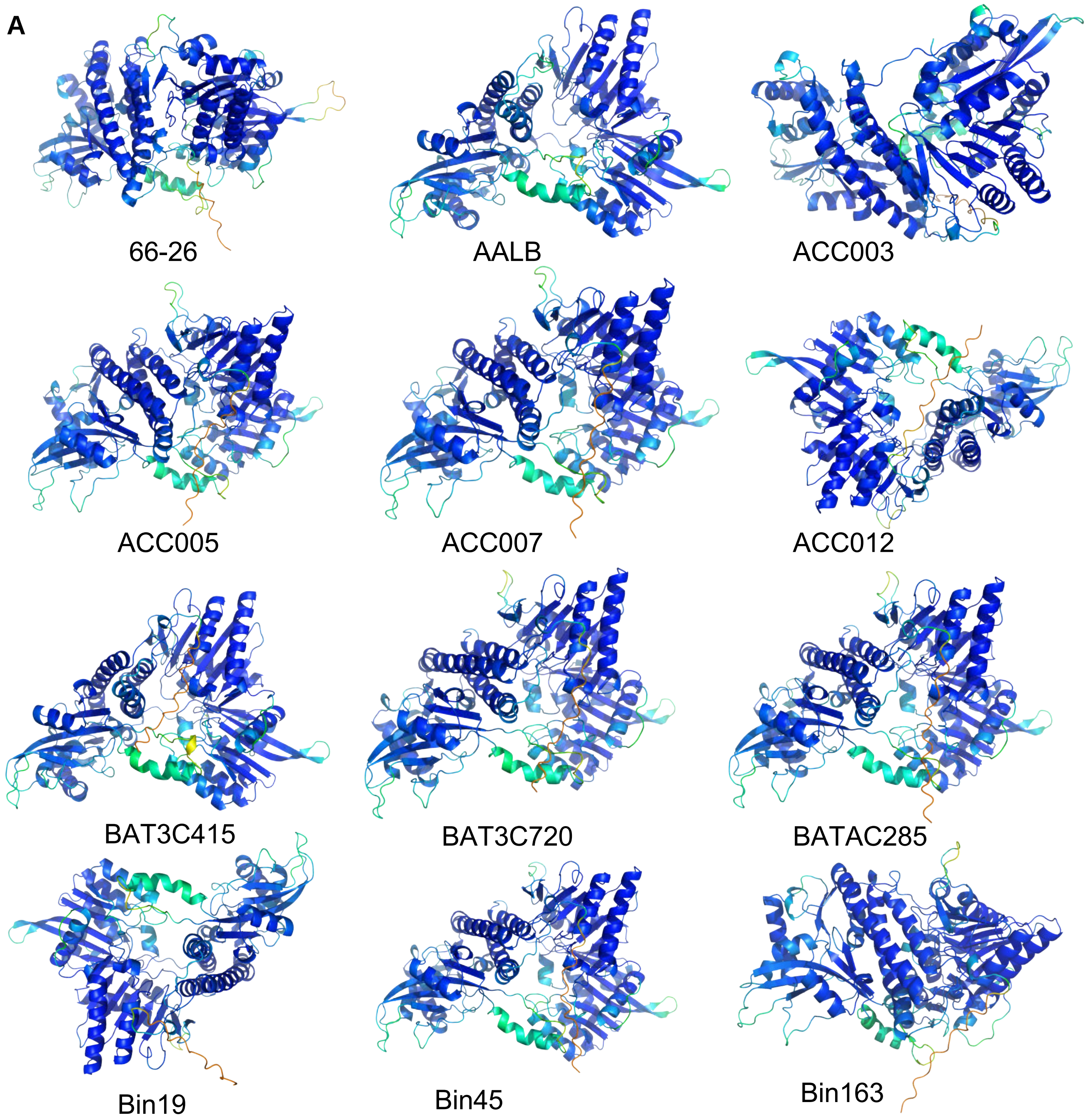
A
66-26
AALB
ACC003
ACC005
ACC007
ACC012
BAT3C415
BAT3C720
BATAC285
Bin19
Bin45
Bin163

**A**

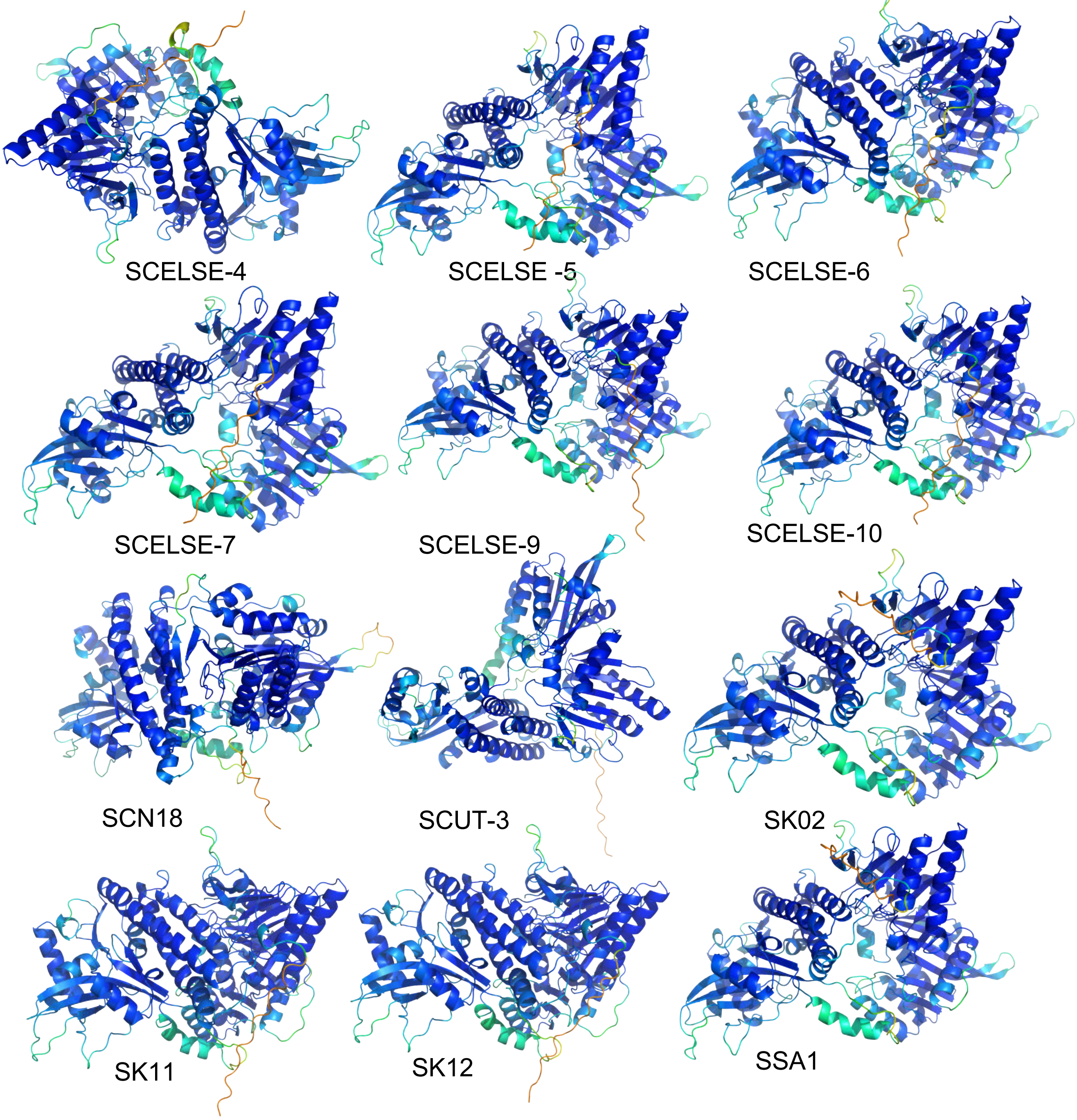

SCELSE-4
SCELSE -5
SCELSE-6
SCELSE-7
SCELSE-9
SCELSE-10
SCN18
SCUT-3
SK02
SK11
SK12
SSA1

**A**

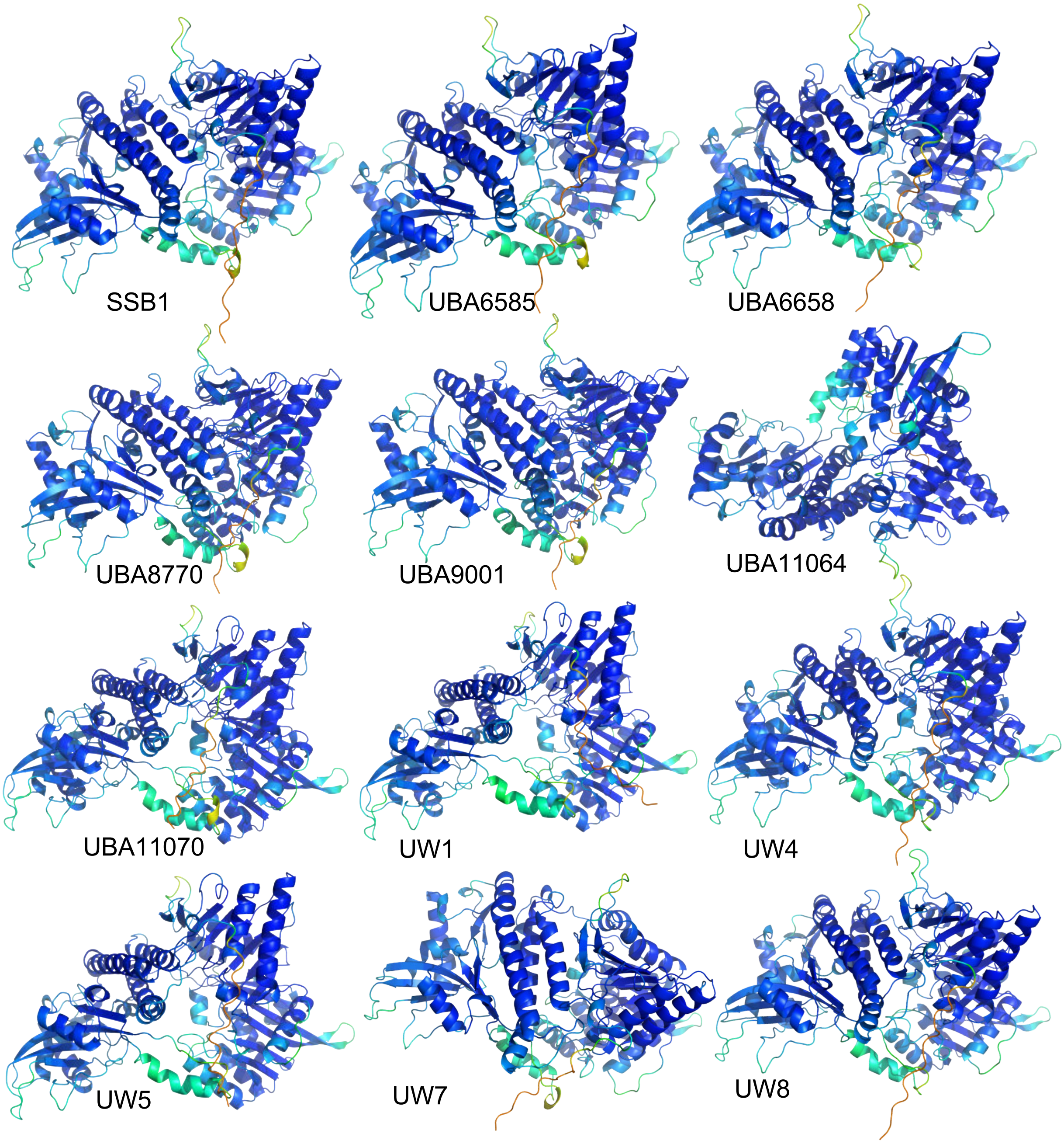
SSB1
UBA6585
UBA6658
UBA8770
UBA9001
UBA11064
UBA11070
UW1
UW4
UW5
UW7
UW8

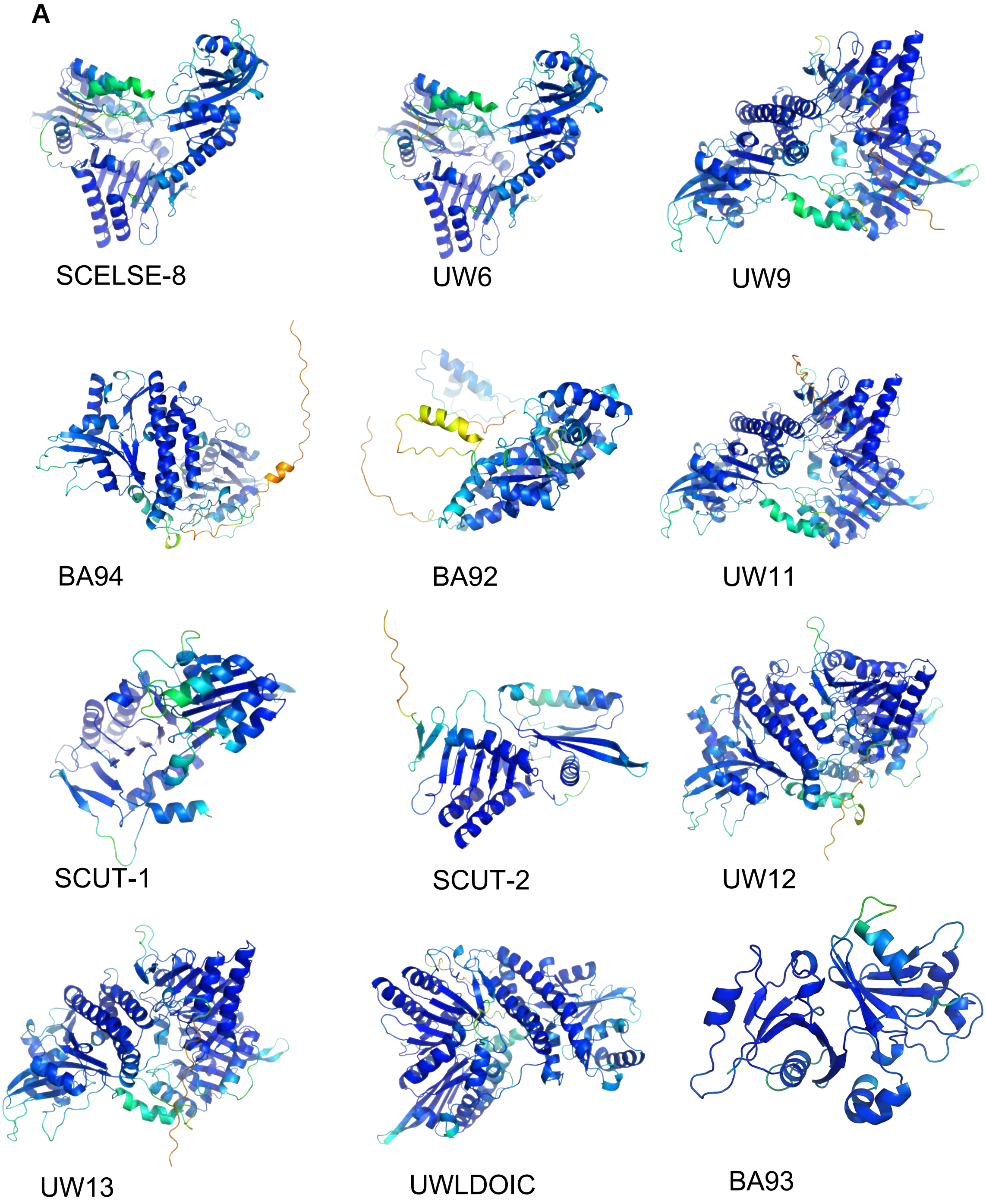
A
SCELSE-8
UW6
UW9
BA94
BA92
UW11
SCUT-1
SCUT-2
UW12
UW13
UWLDOIC
BA93

**B**

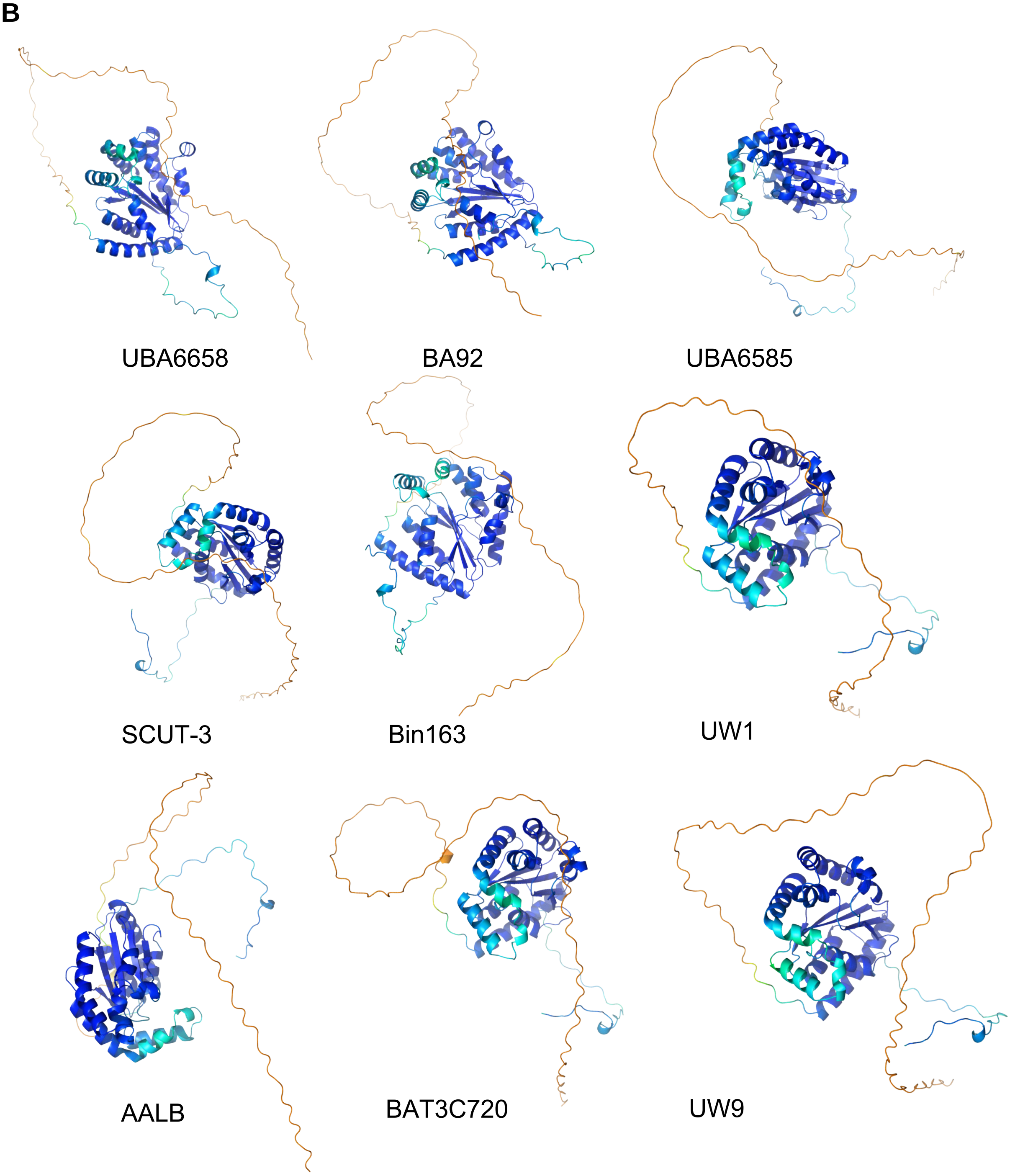

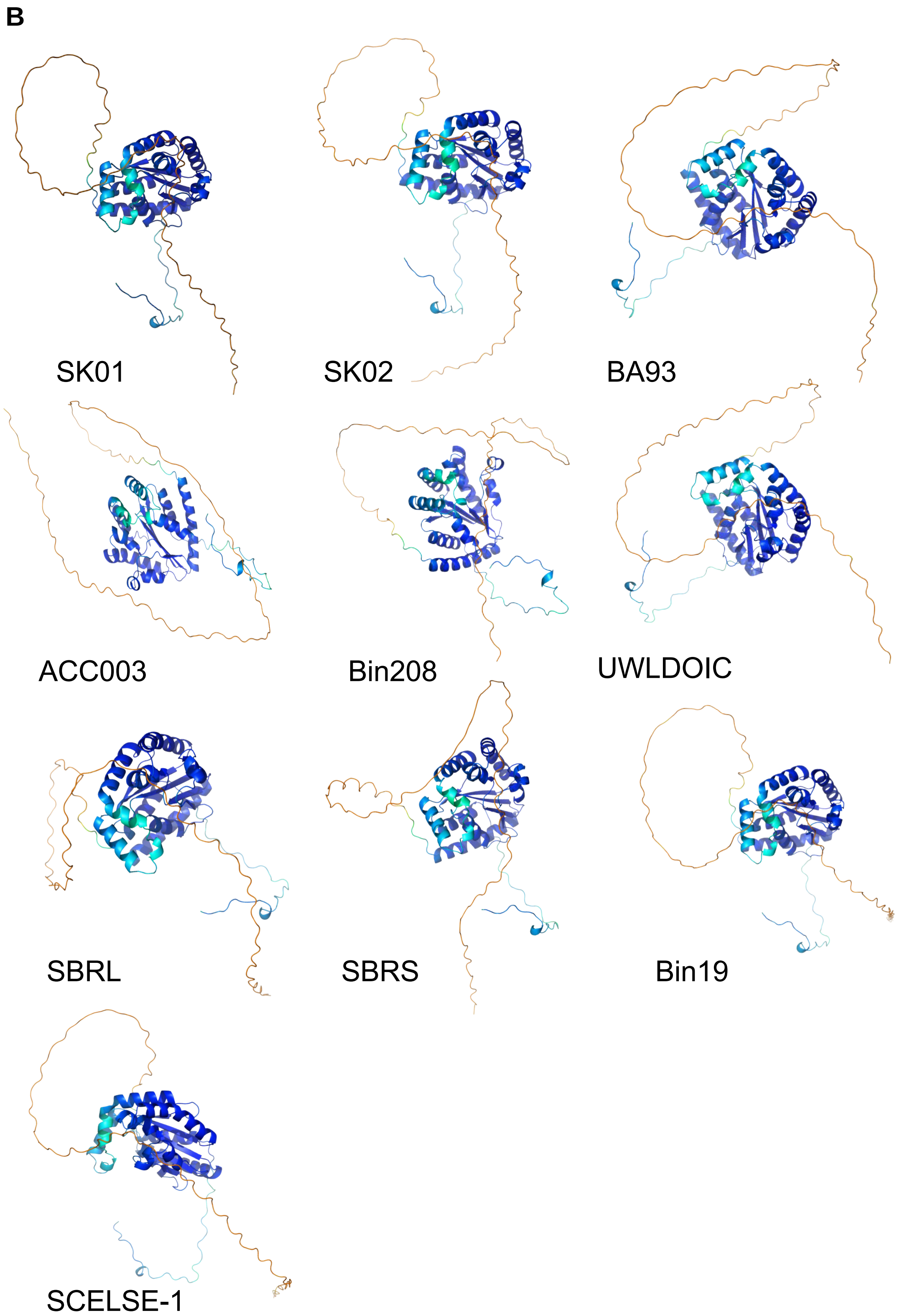
B
SK01
SK02
BA93
ACC003
Bin208
UWLDOIC
SBRL
SBRS
Bin19
SCELSE-1

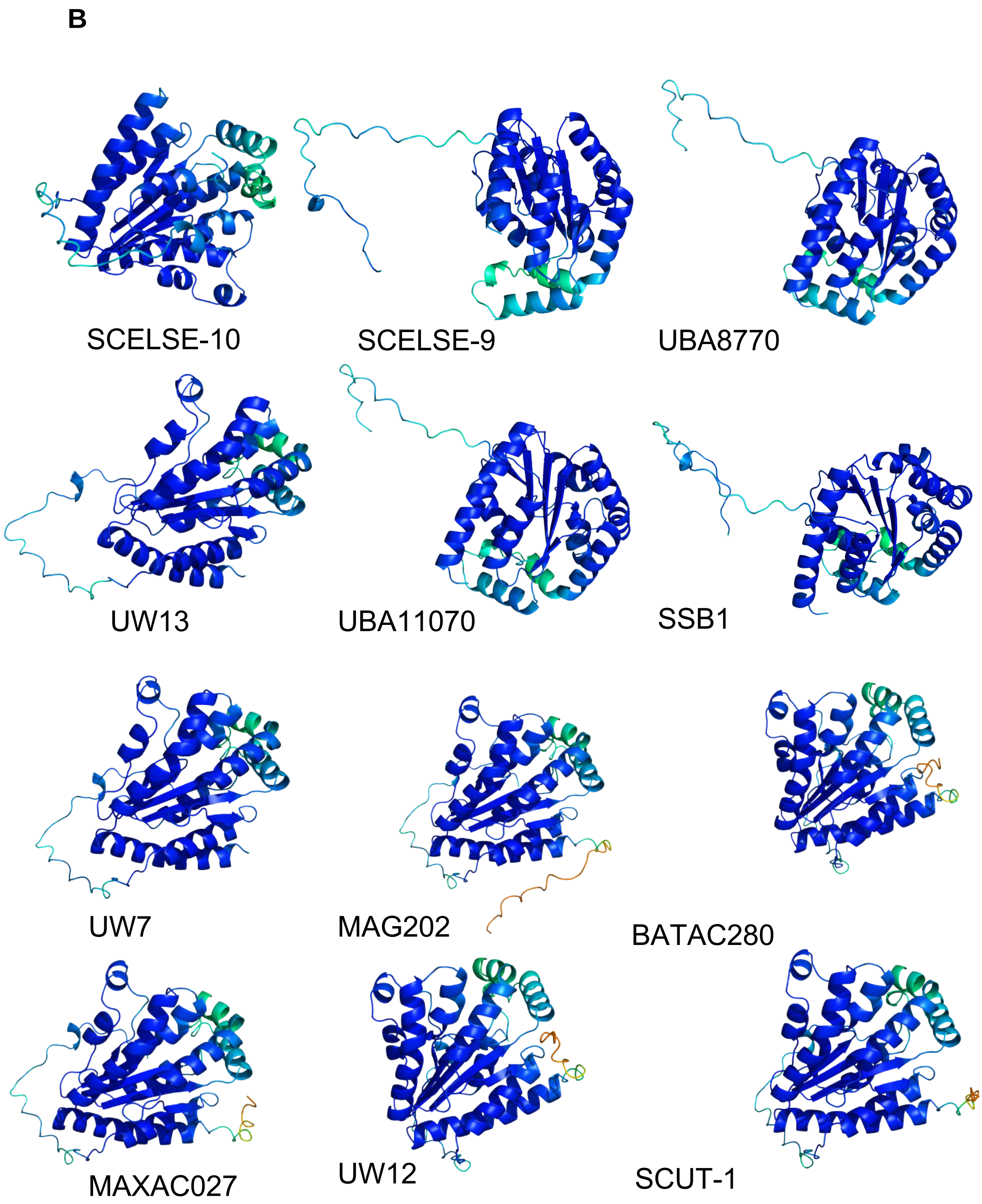
B
SCELSE-10
SCELSE-9
UBA8770
UW13
UBA11070
SSB1
UW7
MAG202
BATAC280
MAXAC027
UW12
SCUT-1

**B**

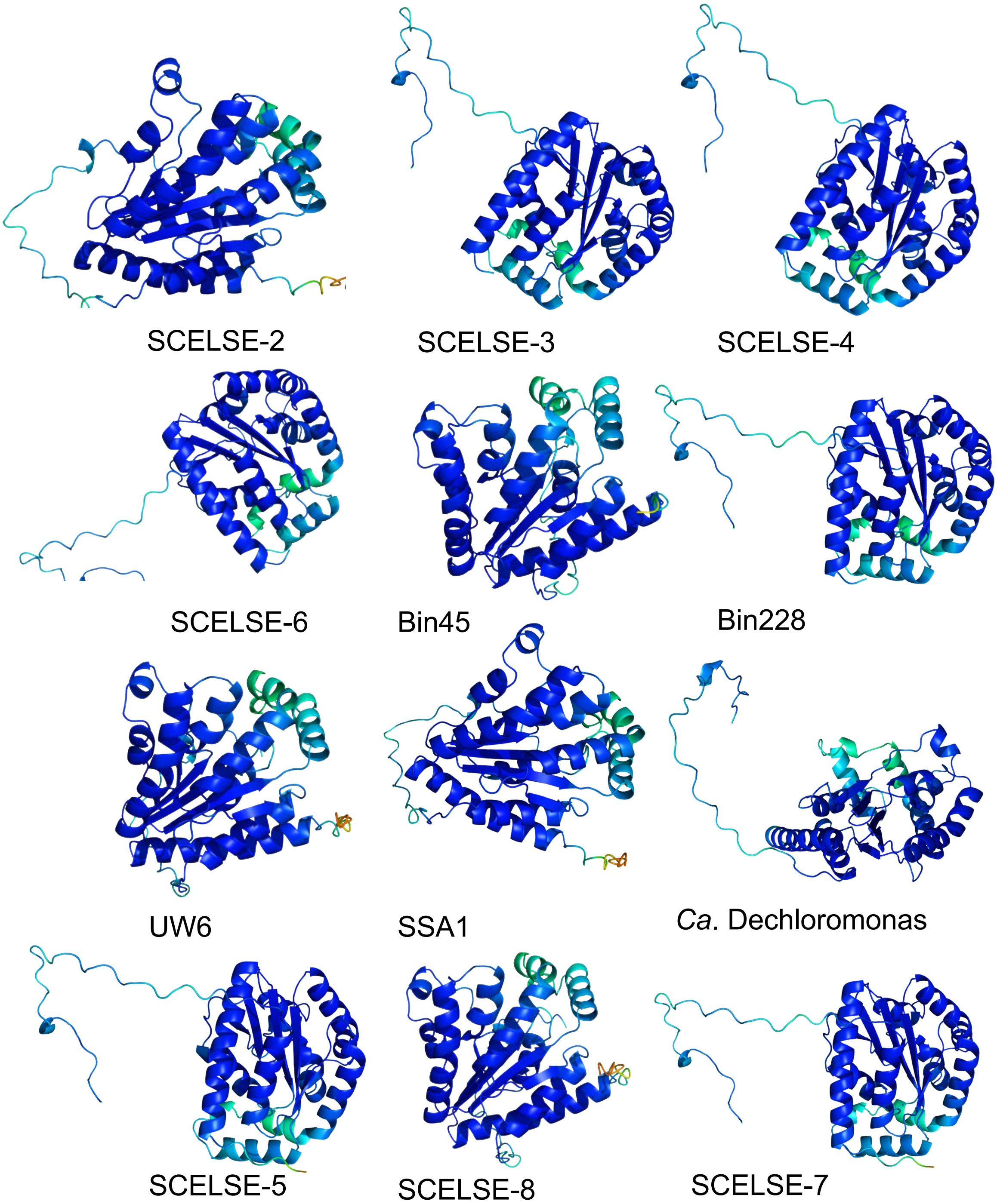

SCELSE-2
SCELSE-3
SCELSE-4
SCELSE-6
Bin45
Bin228
UW6
SSA1
*Ca*. Dechloromonas
SCELSE-5
SCELSE-8
SCELSE-7

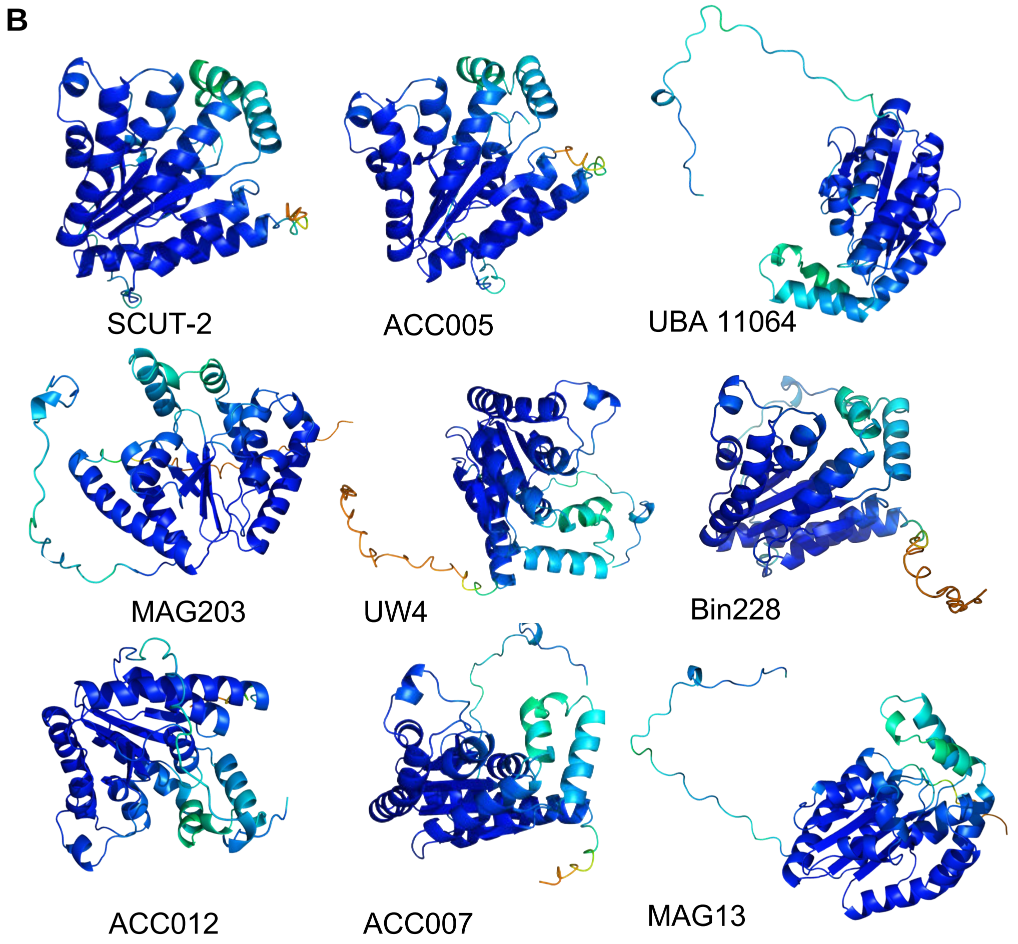
B
SCUT-2
ACC005
UBA 11064
MAG203
UW4
Bin228
ACC012
ACC007
MAG13

**C**

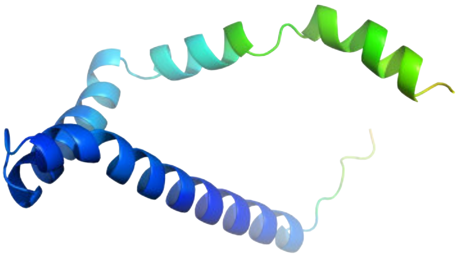

IPGEAEMO_03601(Pit-1)

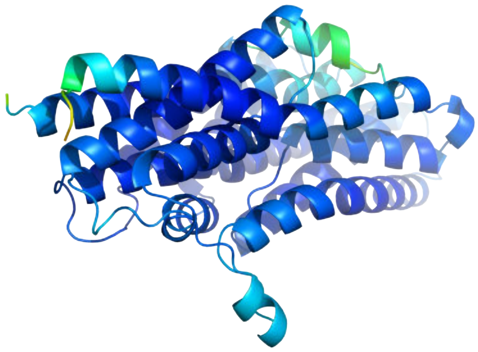

IPGEAEMO_03180(Pit-2)

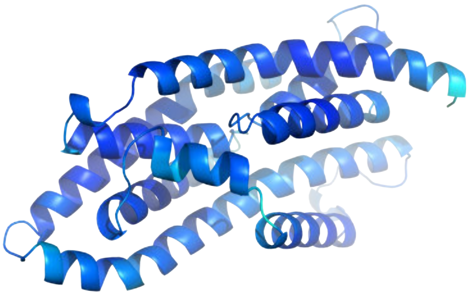

IPGEAEMO_03601(Pit-3)

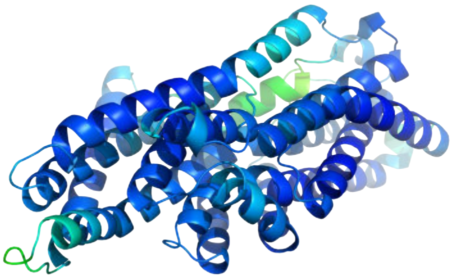

IPGEAEMO_03382(Pit-4)

C

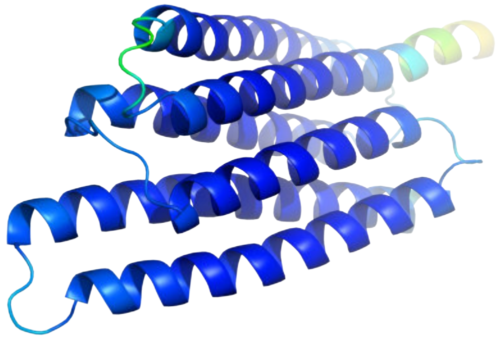

IPGEAEMO_03602(homolog of PhoU-1 )

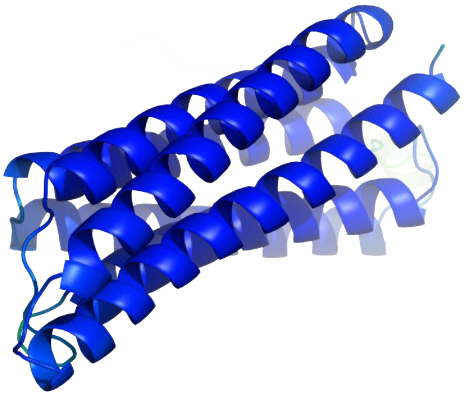

IPGEAEMO_03181(homolog of PhoU-2)

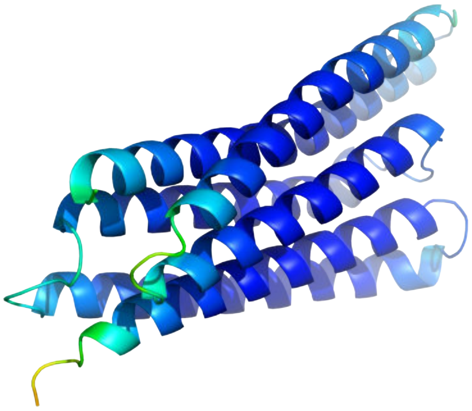

IPGEAEMO_02142(homolog of PhoU-3)

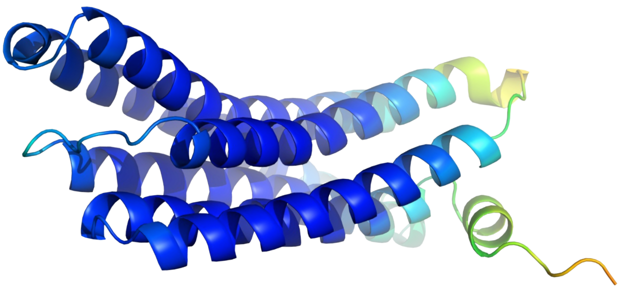

IPGEAEMO_02142(homolog of PhoU-4)

c

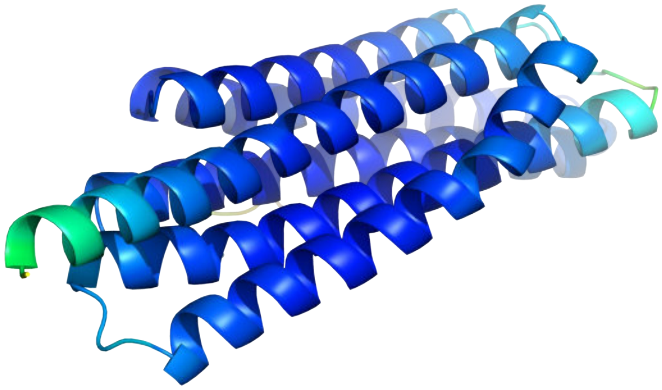

IPGEAEMO_03993(PhoU)

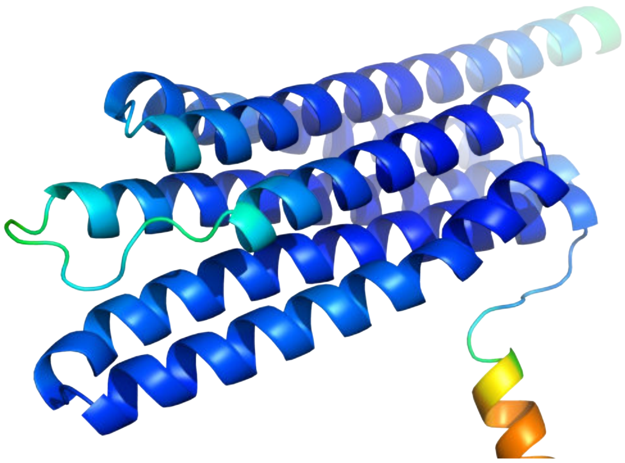

IPGEAEMO_03998(PhoU)

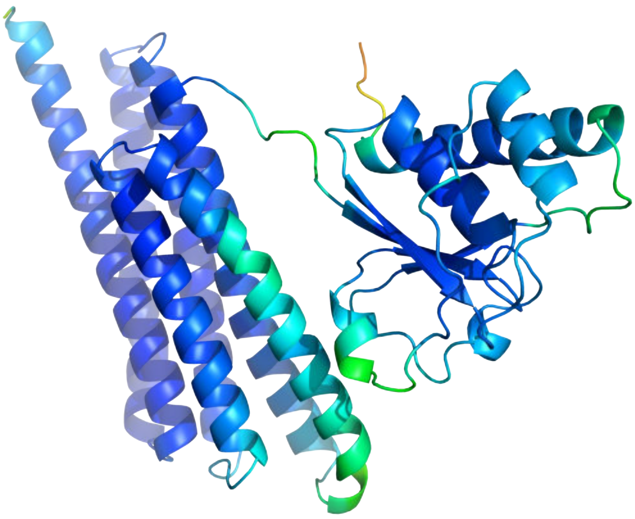

IPGEAEMO_02977 (PhoU)

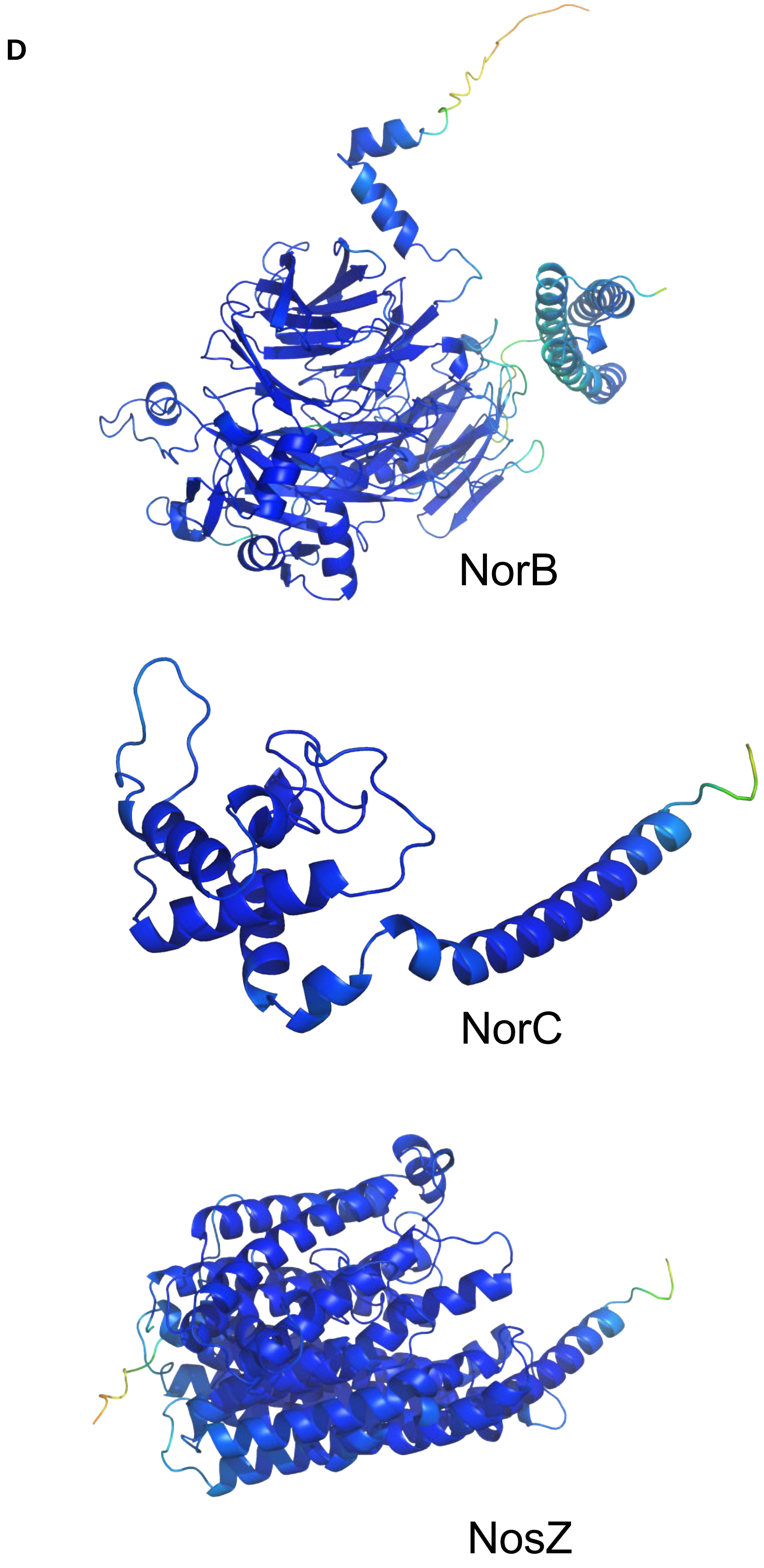
D
NorB
NorC
NosZ

**Figure S2.** The 3D structures of proteins in key (carbon, phosphorus and nitrogen) metabolic pathways of *Ca*. Accumulibacter as predicted by Alphafold2. For each prediction, five models were generated, ranked from 1 to 5, with 1 being the highest pLDDT score. The results obtained using five different models are generally consistent, the model with the highest pLDDT was used for further analyses. Pymol was used to visualize the protein structure, with parameter- [spectrum b, orange_yellow_green_cyan_blue, minimum=40, maximum=100].
(**A**) The structures of PPK1**.** (**B**) The structures of PPK2. (**C**) The structures of phosphate metabolism-related proteins (Pit, PhoU and PhoU homolog). (**D**) The structure of nitrogen metabolism-related proteins (NorC, NorB and NosZ)

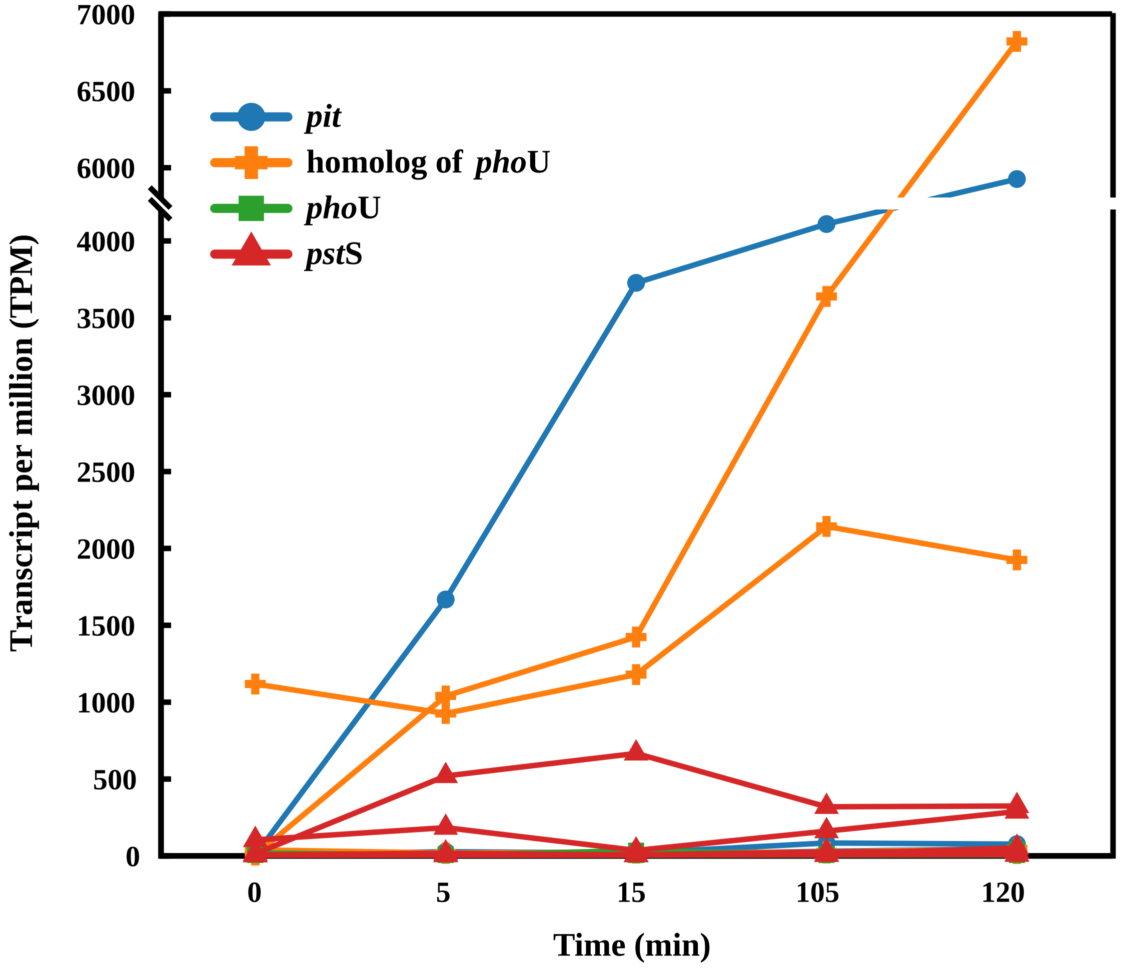


**Figuere S3** Transcriptional dynamics of phosphorus metabolism-related genes in SCUT-2 including *pst*S, *pho*U, *pit* and the homolog of *pho*U.

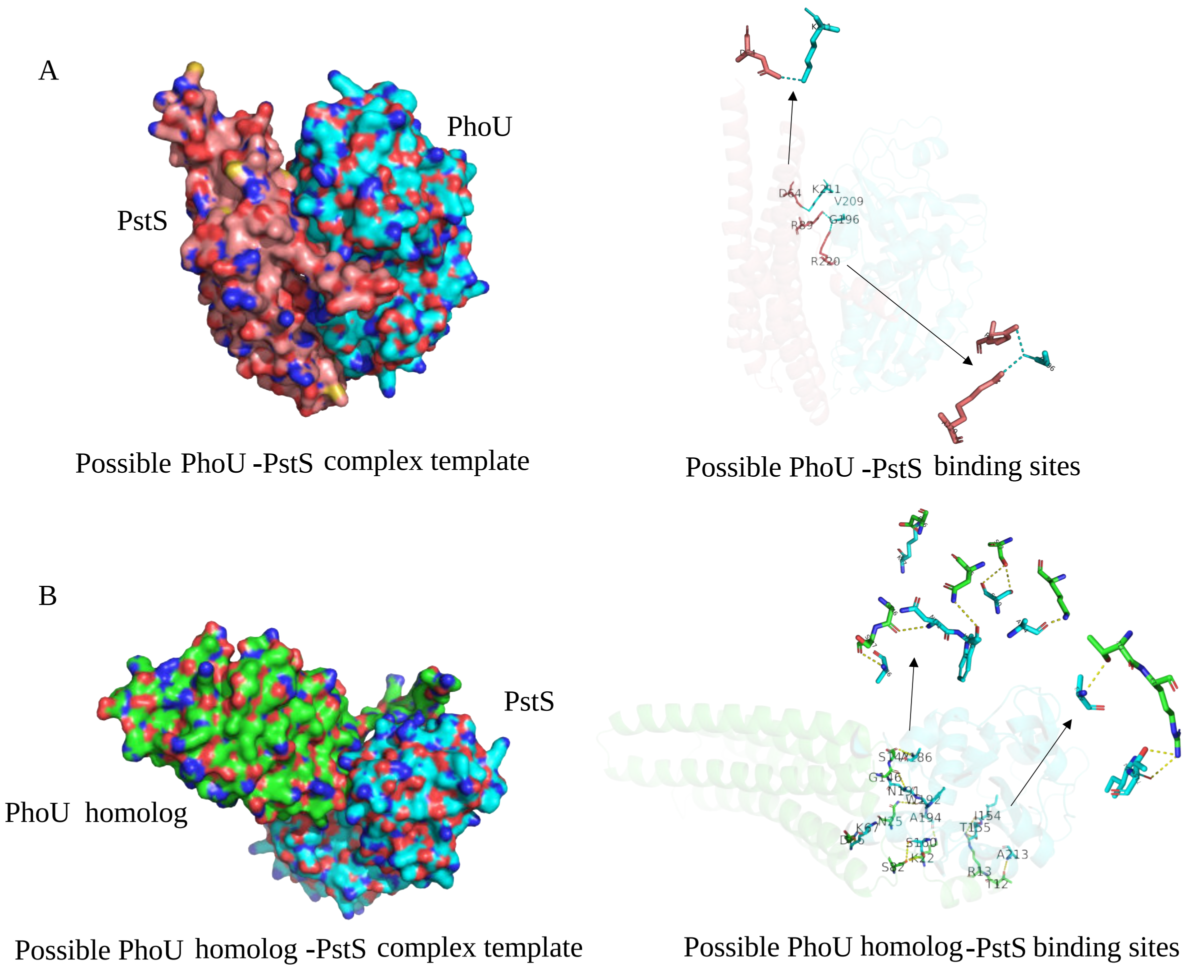


**Figure S4** Phosphorus metabolism-related protein binding models and binding sites predicted by Hdock. Protein-protein docking was visualized using PyMOL. (A)PhoU and PstS binding models and binding sites. (B) PhoU homolog and PstS binding models and binding sites.

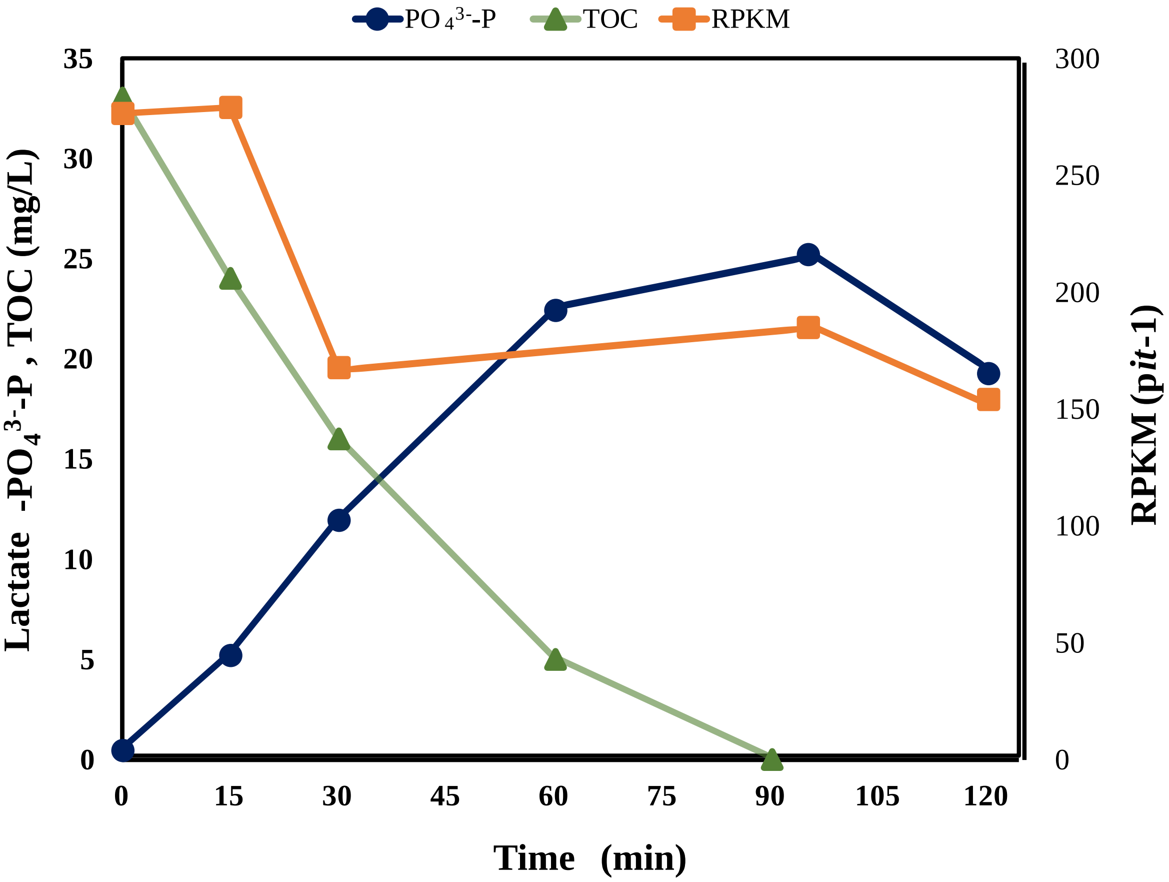


**Figure S5** Full-cycle studies and dynamic gene (*pit*-1) transcription characteristics in SCUT-3. The first 80 min was an anaerobic phase followed by an aerobic phase from 80 min onward. The transit was indicated by a vertical dashed line. RPKM: reads per kilobase of transcripts per million mapped reads.

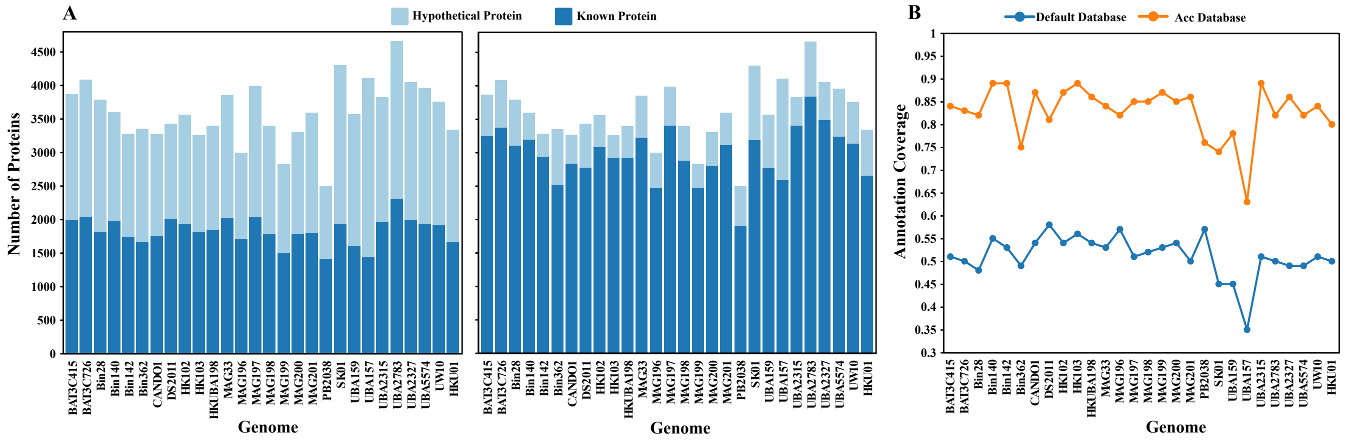


**Figure S6** The efficiency of annotation using the default database and the "Accdata" (a reference database built in this study). (A) The number of hypothetical and known proteins in 28 *Ca*. Accumulibacter based on Prokka using default database (left) and the "Accdata" database. (B) Annotation coverage using the default database (blue) and the "Accdata" database (orange). The latter showed significantly improved coverage.

*Supplementary Spreadsheet*

**Spreadsheet 1**

Typing results based on gene sequences and protein structures. Sheet 1- Pan Rhodocyclaceae gene types identified and the number of genes in each homologous gene type. Sheet 2- Pan Rhodocyclaceae gene types identified and the gene ID in each homologous gene type. Sheet 3- Pan Rhodocyclaceae protein structure types (identified using Foldseek including 17 *Ca*. Accumulibacter genomes from five clades obtained via our scripts by matching the Uniprot ID to the pangenome results and *Ca*. Propionivibrio aalborgensis S1) and the gene ID in each structure type. Sheet 4- A comparison of structure-based typing and sequence-based typing in *Ca*. Accumulibacter similis SCELSE-1 with typing states: These protein pairs were categorized into four groups based on their similarity or dissimilarity in functions, structures, and sequences: (1) non-orthologous protein sequences but with similar structures; (2) orthologous sequences that are structurally different; (3) orthologous sequences with similar structures; and, (4) non-orthologous sequences with different structures. Sheet 5- The result of structural typing between *Ca*. Accumulibacter similis SCELSE-1 and *Ca*. Propionivibrio aalborgensis S1. Sheet 6- The result of sequence-based typing between *Ca*. Accumulibacter similis SCELSE-1 and *Ca.* Propionivibrio aalborgensis S1.

**Spreadsheet 2**

Functional annotation results based on protein structures from 17 *Ca*. Accumulibacter. The molecular function (MF) and enzyme commission (EC) descriptions of all proteins (60,198) were predicted using DeepFRI, generating probability scores within the range of 0 to 1, with each protein assigned at least one gene ontology (GO) term or EC number. The quality of DeepFRI predictions were defined as high quality with scores>0.5 and standard quality with scores >0.2.

**Spreadsheet 3**

*Ca.* Accumulibacter cognatus SCUT-3 and SCUT-2 metatranscriptome. Sheet 1- Reads per Kilobase per million mapped reads (RPKM) of *Ca.* Accumulibacter cognatus SCUT-3 including functions, nucleotide_sequence, etc. Sheet 2 - RAST annotation results for *Ca.* Accumulibacter cognatus SCUT-2, including functions, nucleotide_sequence, etc.

[11]. Arkin AP, Cottingham RW, Henry CS, Harris NL, Stevens RL, Maslov S, Dehal P, Ware D, Perez F, Canon S, Sneddon MW, Henderson ML, Riehl WJ, Murphy-Olson D, Chan SY, Kamimura RT, Kumari S, Drake MM, Brettin TS, Glass EM, Chivian D, Gunter D, Weston DJ, Allen BH, Baumohl J, Best AA, Bowen B, Brenner SE, Bun CC, Chandonia J-M, Chia J-M, Colasanti R, Conrad N, Davis JJ, Davison BH, DeJongh M, Devoid S, Dietrich E, Dubchak I, Edirisinghe JN, Fang G, Faria JP, Frybarger PM, Gerlach W, Gerstein M, Greiner A, Gurtowski J, Haun HL, He F, Jain R, Joachimiak MP, Keegan KP, Kondo S, Kumar V, Land

ML, Meyer F, Mills M, Novichkov PS, Oh T, Olsen GJ, Olson R, Parrello B, Pasternak S, Pearson E, Poon SS, Price GA, Ramakrishnan S, Ranjan P, Ronald PC, Schatz MC, Seaver SMD, Shukla M, Sutormin RA, Syed MH, Thomason J, Tintle NL, Wang D, Xia F, Yoo H, Yoo S, Yu D: **KBase: The United States Department of Energy Systems Biology Knowledgebase**. *Nature Biotechnology* 2018, **36**(7):566-569.

[12]. Kopylova E, Noé L, Touzet H: **SortMeRNA: fast and accurate filtering of ribosomal RNAs in metatranscriptomic data**. *Bioinformatics* 2012, **28**(24):3211-3217.

[13]. van Kempen M, Kim SS, Tumescheit C, Mirdita M, Lee J, Gilchrist CLM, Söding J, Steinegger M: **Fast and accurate protein structure search with Foldseek**. *Nature Biotechnology* 2023.
